# Investigation of reversible histone acetylation and dynamics in gene expression regulation using 3D liver spheroid model

**DOI:** 10.1101/2022.09.22.509080

**Authors:** Stephanie Stransky, Ronald Cutler, Jennifer Aguilan, Edward Nieves, Simone Sidoli

**Author notes:** Corresponding author: Address: 1300 Morris Park Avenue, Bronx, NY 10461, USA.

## Abstract

**Background:** Three-dimensional (3D) cell culture has emerged as an alternative approach to 2D flat culture to model more accurately the phenotype of solid tissue in laboratories. Culturing cells in 3D more precisely recapitulates physiological conditions of tissues, as these cells reduce activities related to proliferation, focusing their energy consumption towards metabolism and homeostasis.

**Results:** Here, we demonstrate that 3D liver spheroids are a suitable system to model chromatin dynamics and response to epigenetics inhibitors. To delay necrotic tissue formation despite proliferation arrest, we utilize rotating bioreactors that apply active media diffusion and low shearing forces. We demonstrate that the proteome and the metabolome of our model resemble typical liver functions. We prove that spheroids respond to sodium butyrate (NaBut) treatment, an inhibitor of histone deacetylases (HDACi), by upregulating histone acetylations and transcriptional activation. As expected, NaBut treatment impaired specific cellular functions, including the energy metabolism. More importantly, we demonstrate that spheroids reestablish their original proteome and transcriptome, including pre-treatment levels of histone acetylation, metabolism, and protein expression once the standard culture condition is restored after treatment. Given the slow replication rate (>40 days) of cells in 3D spheroids, our model enables to monitor the recovery of approximately the same cells that underwent treatment, demonstrating that NaBut does not have long-lasting effects on histone acetylation and gene expression. These results suggest that histone acetylation has minimal epigenetics memory in our spheroids culture.

**Conclusion:** Together, we established an innovative cell culture system that can be used to model anomalously decondensing chromatin in physiological cell growth and rule out epigenetics inheritance if cells recover the original phenotype after treatment. The transient epigenetics effects demonstrated here highlights the relevance of using a 3D culture model system that could be very useful in studies requiring long term drug treatment conditions that would not be possible using a 2D cell monolayer system.

## Background

Long-term regulation of gene expression is a critical driver of epigenetics inheritance, development, and cell memory. Cell cultures are the most common methods to model human biology, including to investigate response to cell stimuli. However, discriminating between short- and long-term responses, e.g. a drug treatment, is not trivial by using cell culture models. This is because cells in culture tend to replicate very fast – on the average every 24-72 hours – meaning that a few days are sufficient to replace most of the original cells that underwent the stimulus. Three-dimensional (3D) cell cultures are a promising *in vitro* alternative to flat cultures (2D) for modeling slow-growing tissue molecular function, diseases, and potential drug penetration. 3D cell cultures create an environment in which multicellular structures grow in any direction interacting with its surroundings in all three dimensions [1], therefore replicating at a slower pace due to the confluency of cells within the 3D structure. Successful 3D culture techniques aim to minimize the formation of necrotic cells while modeling the 3D architecture of parental tissue, as well as recreating a more similar gene expression, signaling, and metabolism of *in vivo* quiescent cells [2–4].

There are a number of 3D cell culturing techniques, which are divided into static or dynamic approaches depending on whether they allow for media exchange. Static 3D cell culturing relies on structural support for cell growth via microplates or through techniques like the “hanging droplet” [5]. These methods allow for high throughput screening, but cells can only be grown into these devices for relatively short amount of time. The dynamic 3D systems allow for the growth of cell aggregates in suspension for virtually an indefinite amount of time thanks to the exchange of cell nutrients and gas [6]. Currently, the most common 3D spheroids are cultured using a scaffold/matrix technique or in a scaffold-free manner, which is very effective but not as high throughput due to laborious cell collection [7]. 3D cell incubators based on the clinostat principle have emerged as alternatives for dynamic systems. The pioneer of this approach is the

Rotary Cell Culture System from Synthecon [8], from which other products such as the ClinoStar from CelVivo were inspired from [9, 10]. The clinostat incubators generate an environment with reduced gravity, which facilitates the formation of highly reproducible 3D structures because organoids and spheroids are assembled by physical properties rather than their natural biological growth [11]. More specifically, there is no net direction for the gravitational effects felt by the spheroids, resulting in a long stable plateau growth phase that better mimics *in vivo* conditions. In addition, cellular proliferation in this 3D system is much more similar to parental tissue, as the doubling time slows from 1 day to 60 days after culturing for 40 days, whereas 2D systems having a doubling time of every 2-4 days [9].

It was proved that clinostat-based cell growth of HepG2/C3A cell lines showed cell phenotypes physiologically closer to that of the *in vivo* liver environment, with comparable levels of ATP, urea, and cholesterol [9, 10]. At the nuclear level, Tvardovsky *et al*. demonstrated that some aspects of chromatin are more similar between primary livers and 3D spheroids compared to 2D cultures; specifically, they used tandem mass spectrometry to study histone proteolytic processing and showed that histones H2B and H3 underwent “clipping” in 3D cells and primary liver but not in 2D cultured cells [12]. This was expected, as 3D cells are characterized by a slower replication rate, enhancing the chromatin properties of a solid tissue. Fast replicating cells rapidly produce new histones, diluting the presence of histone post-translational modifications (PTMs) with slower turnovers [13]. Different studies have also shown the 3D microenvironment as a factor in epigenetic alterations. Feist *et al*. demonstrated that tumor spheroids present a different histone PTMs profile compared to cells growing as a monolayer.

Specifically, they demonstrated that H3K9ac, H3K9me3 and H3K27me3 were differently regulated between the 2D and 3D cultures [14]. Li *et al*. identified thousands of 3D-growth-specific TADs (Topologically Associating Domains) and looping genes and showed that chromatin architecture experience important changes during the 3D culture of breast cancer cells [15].

In this paper, we utilize HepG2/C3A cells to establish a 3D culture system and demonstrate that spheroids can be utilized to investigate the response to epigenetics drugs using RNA-seq, proteomics and selected analysis of metabolites. We show that the chromatin dynamics can be manipulated by drug treatment leading to anomalously accessible chromatin and the effects on gene expression and the proteome can be reversed by restoring the standard culture condition. Altogether, our established 3D liver model is a relatively high throughput platform to investigate long-term effects of drug treatment on gene expression in human cells.

## Results

### Generation of functional spheroids using HepG2/C3A cells

To obtain liver spheroids from HepG2/C3A cells, we optimized the protocol originally described by Wrzesinski and colleagues [9] for ease of use and to achieve reproducible and cost-effective culturing of >100 spheroids per bioreactor. In brief, the protocol initiated by utilizing HepG2/C3A flat culture (2D culture) grown until reached 80% confluency (**Fig. 1A**). To culture cells as spheroids (3D culture), 1.2×10^6^ cells were plated in an ultra-low attachment plate containing microwells (approximately 2,000 cells per microwell) and were incubated for 24 h to allow them to self-aggregate (**Fig. 1B**). Before transfer to a bioreactor, the size and roundness of spheroids were analyzed by microscopy to ensure a uniform culture. Newly formed HepG2/C3A spheroids (**Fig. 1C**) ranged from 152 to 232 µm (**Fig. S1A)**, with an average size of 201 µm (± 18.3 µm). As spheroids get old, they get bigger and present a denser core compared to the borders (**Fig. 1D** and **1E**). Although they are morphologically formed after 2 weeks, they only get functionally active after 3 weeks in culture, as previously demonstrated [10]. The spheroid shown in **Fig. 1F**, which was maintained in culture for 22 days, represents a functionally mature spheroid.

**Fig. 1.**
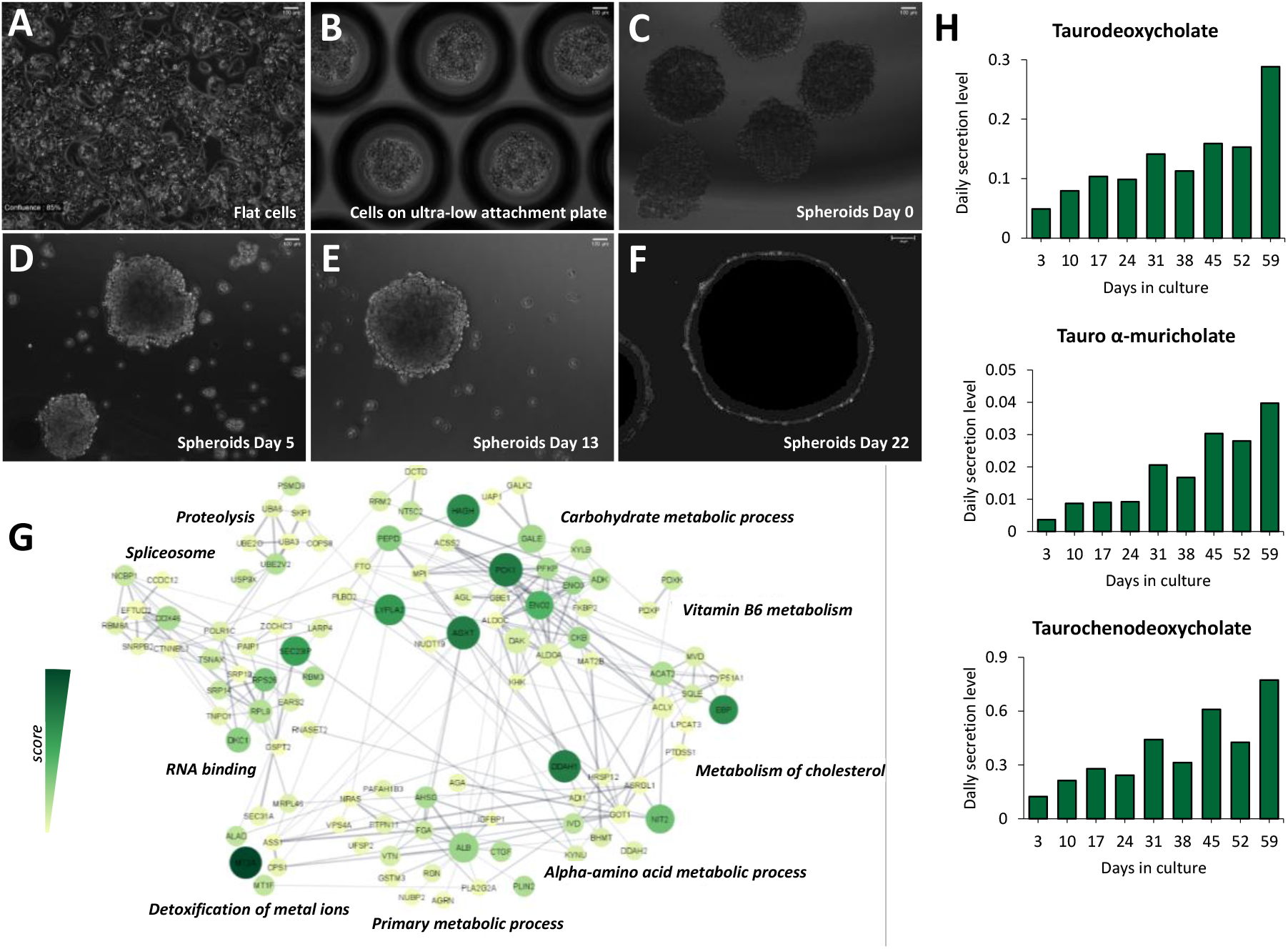
The proteome and metabolome of HepG2/C3A spheroids resemble a functional liver. **(A)** HepG2/C3A (human hepatocellular carcinoma) flat cells were cultivated in DMEM media supplemented with 10% FBS, until colonies reached 80% confluence. **(B)** Then, cells were trypsinized and plated on an ultra-low attachment 24 well plate containing microwells. **(C)** Spheroids were then detached from the plate, transferred to a bioreactor and cultivated in the same media as flat cells (day 0). **(D)** Spheroids on day 5, **(E)** day 13 and **(F)** day 22. Images were acquired using PAULA Smart Cell Imager (Leica) with 10x magnification. Scale bar = 100µm. **(G)** Clustered network of the 200 up-regulated proteins in HepG2/C3A spheroids. Clusters reflect connected portions of the network and correspond to functional categories of the proteins. Size of nodes represents *p-value*, color darkness represents the score (fold change enrichment times the *p-value*) and line thickness represents the score of interaction confidence retrieved from the software String (v11, https://string-db.org). The network was constructed by Cytoscape [18]. **(H)** Bar graphs show the daily secretion levels of the bile salts taurodeoxicholate, tauro α-muricholate, and taurochenodeoxycholate detected in the supernatant of spheroid culture.

By growing HepG2/C3A cells as 3D spheroids, we verified whether the expressed proteome resembled more closely the one of functional liver. Biological processes significantly enriched in HepG2/C3A spheroids revealed proteins mostly related to the carbohydrate, cholesterol, and vitamin B6 metabolisms (**Fig. 1G**). Proteins related to amino acid metabolism and detoxification of metal ions were also enriched in spheroids (**Fig. 1G**). In fact, besides the role in metal detoxification, metallothioneins also contribute to the antioxidant activity and protective effects against free radicals performed by the liver [16]. We identified 5 isoforms of metallothioneins (MT1E, MT1F, MT1G, MT1X, and MT2A), all more abundant in spheroids in comparison to flat cells (**Fig. S1B**). We were also able to detect some biomarkers known to be predominant in the liver, whether in disease or normal conditions, such as alpha-fetoprotein (AFP), albumin (ALB), and apolipoprotein E (APOE), lactate dehydrogenase (LDHA), alanine aminotransferase (GPT2), and aspartate aminotransferase (GOT2) (**Fig. S1C**). It is worth pointing out that our data was log-transformed and normalized to correct for any artificial biases in sample pipetting and injection that consequently could lead to differences in protein content. Therefore, the differences seen in the number of proteins between flat cells and spheroids indicate relative changes of protein expression and not an overall higher content of protein in the spheroids.

We also aimed to demonstrate that the longer spheroids are cultured, the more they produce and secrete metabolites typical of liver functions. Metabolic profiling of collected cell media from spheroid cultures revealed increasing levels of the bile salts taurodeoxycholate, tauro α-muricholate, and taurochenodeoxycholate (**Fig. 1H**), which are known to be produced by the liver. The full metabolite profiling of the cell media is available as **Supplementary Table 1**.

Together, these results confirmed that HepG2/C3A cells grown using the ClinoStar (CelVivo) 3D technology resemble the typical characteristic of functional liver, which includes maintaining whole-body lipid, glucose and energy metabolism, as well as detoxification of drugs, xenobiotics and metals [17].

### Liver spheroids respond to chromatin decondensation

To evaluate the sensitivity of our 3D model to drug treatment, spheroids were maintained in a growth media containing 20 mM of sodium butyrate (NaBut) for 3 days (**Fig. 2A**). After the collection of spheroids and culture supernatant, the treatment media was replaced by the standard growth media, and spheroids were maintained in culture for additional 7 days, to evaluate whether they could recover their normal physiology. Sodium butyrate is known to act as a histone deacetylase inhibitor (HDACi), favoring histone acetylation and thus remodeling of chromatin towards an open and transcriptionally active state [19]. HDACi can also have anti-cancer effects, which depend on the type of cancer and the treatment dose [20]. We analyzed histone modifications by mass spectrometry, using the protocol optimized by Sidoli and Garcia [21]. Obtained chromatograms were extracted using EpiProfile [22].

**Fig. 2.**
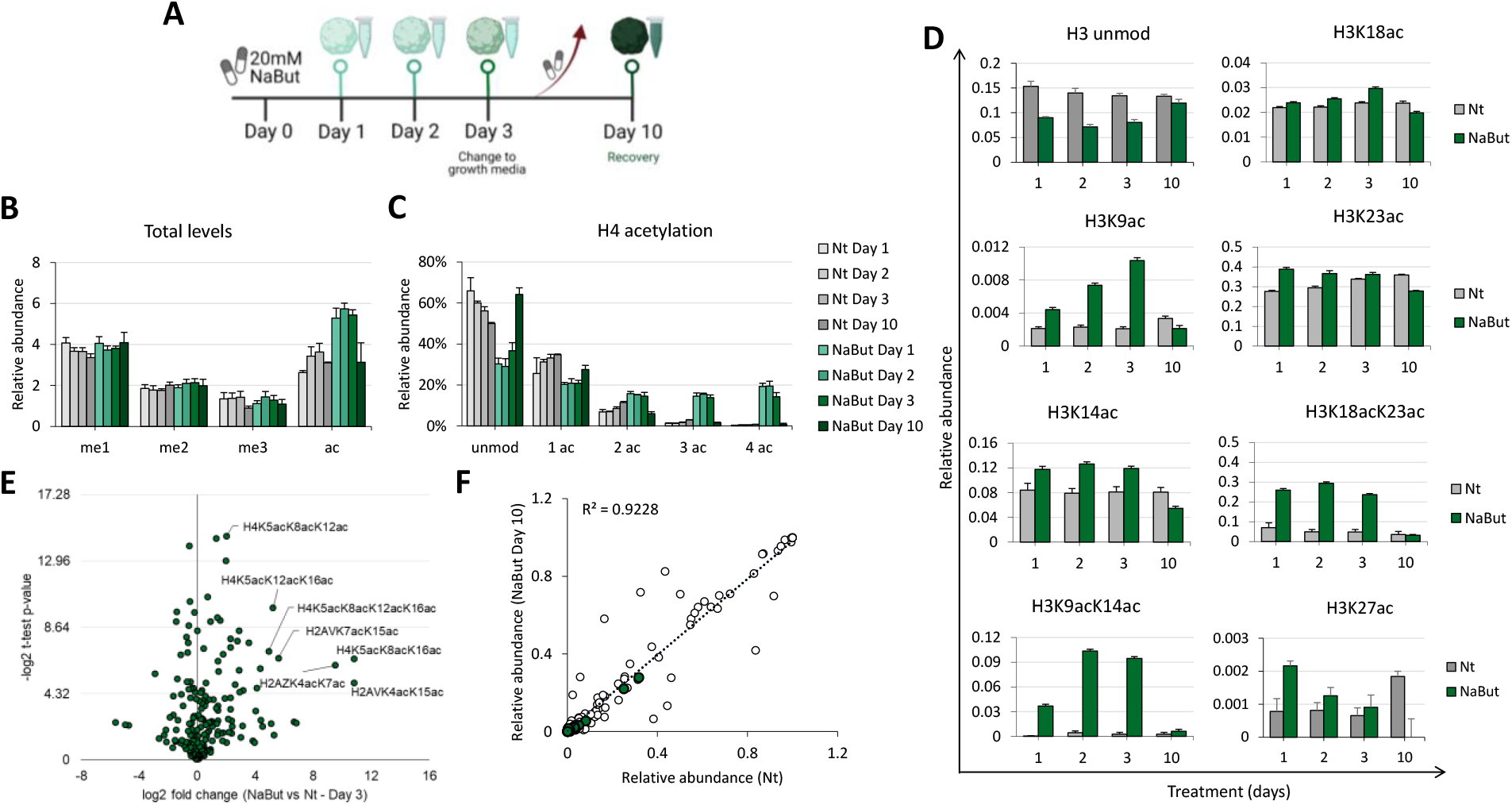
Spheroid respond to chromatin decondensation. Spheroids were treated with 20 mM of NaBut and were kept in culture for 3 days. After collection, the treatment media was replaced by the standard growth media and spheroids were maintained for additional 7 days, until the last collection on day 10. Histones were extracted from spheroids and analyzed by mass spectrometry. **(A)** Workflow for spheroids treatment. **(B)** Total levels of histone peptides containing 1, 2 or 3 methylations (me1, me2, me3, respectively) or containing acetylations (ac). **(C)** Total levels of histone H4 peptides containing acetylations (1ac, 2ac, 3ac, 4ac). Unmod, unmodified peptide. **(D)** Relative abundance of histone H3 acetylated peptides. Data are represented as means ± SD. Nt, non-treated. **(E)** Volcano plot representing NaBut vs Nt fold change after 3 days of treatment and **(F)** Correlation coefficient (R) of histone marks between non-treated (Nt) spheroids and NaBut-treated spheroids after recovery (Day 10). Highlighted green dots correspond to acetylated histone peptides.

Overall, the total levels of mono, di or trimethylated histones (me1, me2, me3, respectively) did not differ between non-treated (Nt) and NaBut-treated groups (**Fig. 2B and Fig. S2A**). However, the acetylation (ac) levels increased after treatment (**Fig. 2B**). By using our protocol, histone H4 is processed in a peptide carrying up to 4 acetyl groups (amino acids 4 to 17, modification sites being K5, K8, K12 and K16). Therefore, we focused on that peptide to evaluate the accumulation of hyperacetylation on histone proteins. Treatment with NaBut for 3 days increased in almost 20% the levels of histone H4 peptides containing multiple acetylations, i.e. 3ac and 4ac (**Fig. 2C**). Hyperacetylation of histones, triggered by NaBut, occurs slower in spheroids compared to flat cells (**Fig. S2B**), which is expected as the lower proliferation rate of cells in the 3D culture leads to slower production of new histones to be modified. Acetylation marks on histone H3 peptides were also quantified, and a similar trend was identified in both spheroids and flat cells (**Fig. 2D and Fig. S2C**, respectively). H3K9ac as well as the combinatorial PTMs H3K9acK14ac and H3K18acK23ac, were the marks most upregulated by the treatment.

Importantly, we demonstrated that after 7 days in a cell culture media with no treatment, spheroids were able to reestablish their normal levels of histone modifications physiology. By determining cell viability (adenylate kinase release), we demonstrated that spheroids were able to recover from the treatment due to the reduced number of damaged cells (**Fig. S3A**). Furthermore, it is evident that on day 10 the levels of histone marks are not significantly different from their levels in the non-treated (Nt) condition (**Fig. 2B-D**). The recovery event can also be seen in **Fig. 2E** and **2F**; the volcano plot clearly demonstrates that spheroids treated for 3 days show an enrichment of histone H4 hyperacetylated peptides (**Fig. 2E**), while on day 10 the levels return to basal levels and show a high correlation (R^2^ = 0.9228) with the non-treated condition (**Fig. 2F**). Besides showing that spheroids recover their normal metabolism and revert the effects caused by NaBut, our results suggest that histone acetylation seems to have minimal memory in our cell culture, as it is not being conserved once treatment has been removed.

It is worth mentioning that this analysis is prohibitive in 2D cell monolayer, as 10 days is a sufficient time frame to fully replicate all the cells in culture, i.e. the recovery analysis would be performed on daughter cells rather than those which received the treatment. This highlights the relevance of the 3D culture model described here. To demonstrate that cells in 3D spheroids have a very slow replication rate, we quantified the relative abundance of the enzyme DNMT1, which is known for its role in maintaining methylation pattern in newly synthesized DNA strand [23]. We demonstrated that the levels of DNMT1 are lower in spheroids compared to flat cells (**Fig. S3B**). Moreover, by metabolic labeling we confirmed that the protein turnover rate is significantly higher in flat cells than in spheroids, i.e. approx. 20% labeling after 2 days vs 1% labeling of proteins in spheroids (**Fig. S3C**). This demonstrates that cells in 3D spheroids do not produce new DNA and new proteins, phenomena commonly associated with cell proliferation.

### Transcriptional alteration and recovery from chromatin decondensation

We had demonstrated that NaBut treatment induced histone hyperacetylation (**Fig. 2B-F**). Since histone acetylation is known to directly and indirectly induce transcriptional activation [24], we aimed to demonstrate whether gene expression is affected by our treatment and it is restored once the treatment is removed. We performed RNA sequencing (RNA-seq) of treated and non-treated spheroids, identifying 6230 genes with significantly different (FDR < 0.05) trajectories due to NaBut treatment (**Fig. 3A**). K-means clustering of this set of genes broke them down into 8 clusters where genes in clusters A-D are initially upregulated and genes in clusters E-G following the opposite pattern over the time-course (**Supplementary Table 2**).

**Fig. 3.**
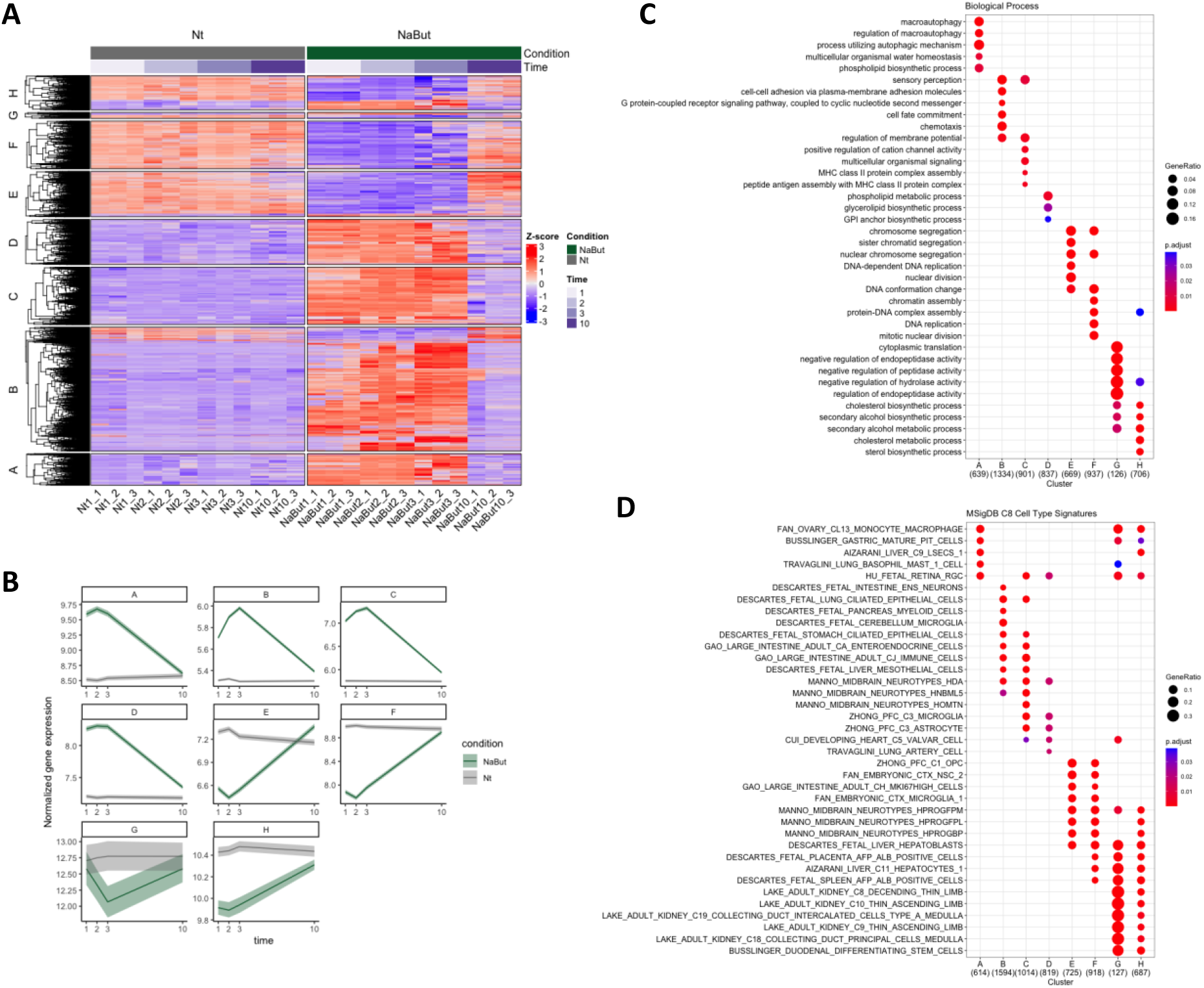
Transcription alteration and recovery following NaBut treatment. **(A)** Heat map showing genes significantly altered over the time-course of the treatment. Control (C1, C2, C3, C10) and NaBut-treated (N1, N2, N3, N10) spheroids, treated for 1, 2 or 3 days and the recovery time point (day 10); n=3. **(B)** Average trajectory of genes over the time-course for each cluster showing return to baseline levels at recovery time point (day 10). Ribbons represents standard error. **(C)** Gene ontology biological process over-representation analysis for each of the clusters. **(D)** Cell type signature over-representation analysis for each of the clusters. Clusters A-D are down regulated and clusters E-H are up regulated over the time-course.

Plotting the data onto a heat map, this clearly showed distinct transcriptional profiles between Nt and NaBut-treated spheroids over the time course (**Fig. 3A**), highlighting that the majority of genes were upregulated in presence of chromatin hyperacetylation. Although the clusters show different trajectories after treatment, nearly all affected genes return to the control levels by day 10 (NaBut10) where the spheroids were allowed to recover for 7 days (**Fig. 3B**). In order to get an insight into the altered biological processes induced by histone hyperacetylation and chromatin decondensation, over-representation analysis for each of the clusters was performed using either the Gene Ontology or Molecular Signatures databases. Biological processes that were impaired by the treatment represented a slowing of cell division and metabolism, such as cholesterol metabolism (cluster G and H), DNA replication related chromatin changes (cluster E and F), and alcohol/sterol metabolism (cluster C) **(Fig. 3C**). On the other hand, a myriad of seemingly unrelated biological processes were enhanced by the treatment which included macroautophagy (cluster A), cell-cell adhesion and chemotaxis (cluster B), calcium activity (cluster C), and phospholipid/glycerolipid biosynthesis (cluster H). This indicates that chromatin decondensation induced by non-specific hyperacetylation led to a spurious upregulation of genes rather than a coordinated response to the treatment. To confirm this, enrichment for cell type signatures showed loss of liver cell identity after treatment, such as down regulation of the hepatocyte and hepatoblasts (clusters E-H) and the up regulation of non-liver signatures such as macrophage (cluster A), neurons (clusters B-C) microglia, astrocytes, heart, and lung (clusters C-D) **(Fig. 3D**).

### Histone hyperacetylation and chromatin decondensation have a direct effect on the spheroids proteome

Given the effect of NaBut on cellular biological processes at the transcriptome level, we then investigated the gene ontology (GO) enrichment at the proteome level (**Fig. 4**). GO analysis of upregulated proteins of spheroids treated with NaBut for 1, 2 and 3 days (**Fig. 4A, B** and **C**, respectively) revealed a poor enrichment with non-specific cellular processes, indicating a similar uncontrolled gene expression as previously detected by RNA-seq data. On the other hand, the top enriched GO terms for the upregulated proteins of spheroids after the recovery period (day 10) revealed liver-related functions, such as lipid metabolic process, cholesterol transport and plasma lipoprotein particle remodeling (**Fig. 4D**). The full list of identified proteins is available as **Supplementary Table 3**.

**Fig. 4.**
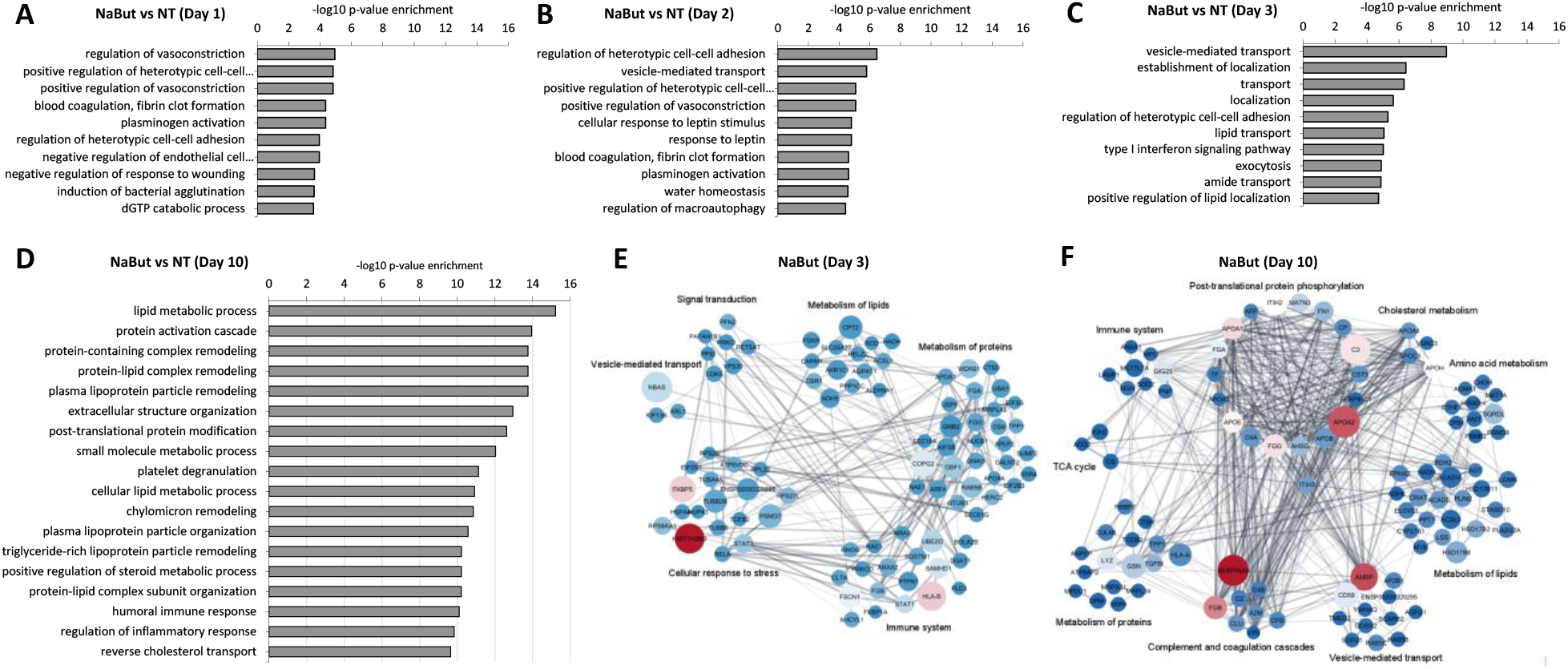
Liver function is impaired by chromatin decondensation and transcription activation. Gene ontology (GO) enrichment analysis of spheroids treated for **(A)** 1, **(B)** 2, **(C)** 3 days and **(D)** after recovery (day 10). Functional annotation was obtained using GOrilla [25]. Clustered network of the 200 up-regulated proteins in spheroids treated with NaBut for 3 days **(E)** and **(F)** recovered spheroids (day 10). Clusters reflect connected portions of the network and correspond to functional categories of the proteins. Size of nodes represents *p-value*, color darkness represents the score (fold change enrichment times the *p-value*) and line thickness represents the score of interaction confidence retrieved from the software String (v11, https://string-db.org). The network was constructed by using Cytoscape [18].

Proteins with the highest score in spheroids treated with NaBut for 3 days or spheroids recovered from treatment (day 10) were then used to construct a network of protein interactions (**Fig. 4E** and **F**, respectively) to demonstrate the molecular and cellular mechanisms enriched in each of the conditions. Biological processes significantly enriched in treated spheroids revealed proteins mostly related to cellular response to stress and immune system, although proteins related to metabolism of lipids and proteins and signal transduction were also enriched (**Fig. 4E**). Interestingly, the pathways most enriched after spheroids are recovered from the treatment include post-translational protein phosphorylation, vesicle-mediated transport, and complement and coagulation cascades (**Fig. 4F**). Biological processes related to liver function such as the TCA cycle, metabolism of lipids and cholesterol metabolism are also enriched once spheroids reestablish their homeostasis, indicating that the cells within 3D spheroids regulate their proteome towards recovering original liver functionalities of cells in 3D spheroids. Interestingly, this regulation was not noticed in RNA-seq data, most likely due to the faster turnover or RNAs compared to proteins. In fact, by day 10 basically all RNA levels were back to baseline (**Fig. 3A**).

Altogether, our data show that NaBut treatment for 3 days induces histone hyperacetylation and, as indirect effects, it triggers transcriptional activation most likely through chromatin decondensation. However, the proteome showed a slower regulation compared to histone modifications and mRNA, potentially contributing to re-establishing the baseline functionalities of pre-treated spheroids. Enriched proteins related to lipid metabolic process, described above, are highlighted in the volcano plots for each of the treatment days (**Fig. 5A**). It is evident that most of these proteins are enriched on day 10, when the cells are no longer exposed to the treatment, indicating that cells returned their normal metabolism. On the other hand, proteins related to response to treatment (**Fig. 5B**) and transcription (**Fig. 5C**) are enriched specially on day 3 of NaBut treatment, suggesting that the effect of NaBut treatment might be cumulative in the cells. Altogether, this indicates that our 3D cell culture is a suitable model for recovery studies as well as long-term treatments (model in **Fig. 6A-B**), approaches that are not supported by cells growing as a monolayer.

**Fig. 5.**
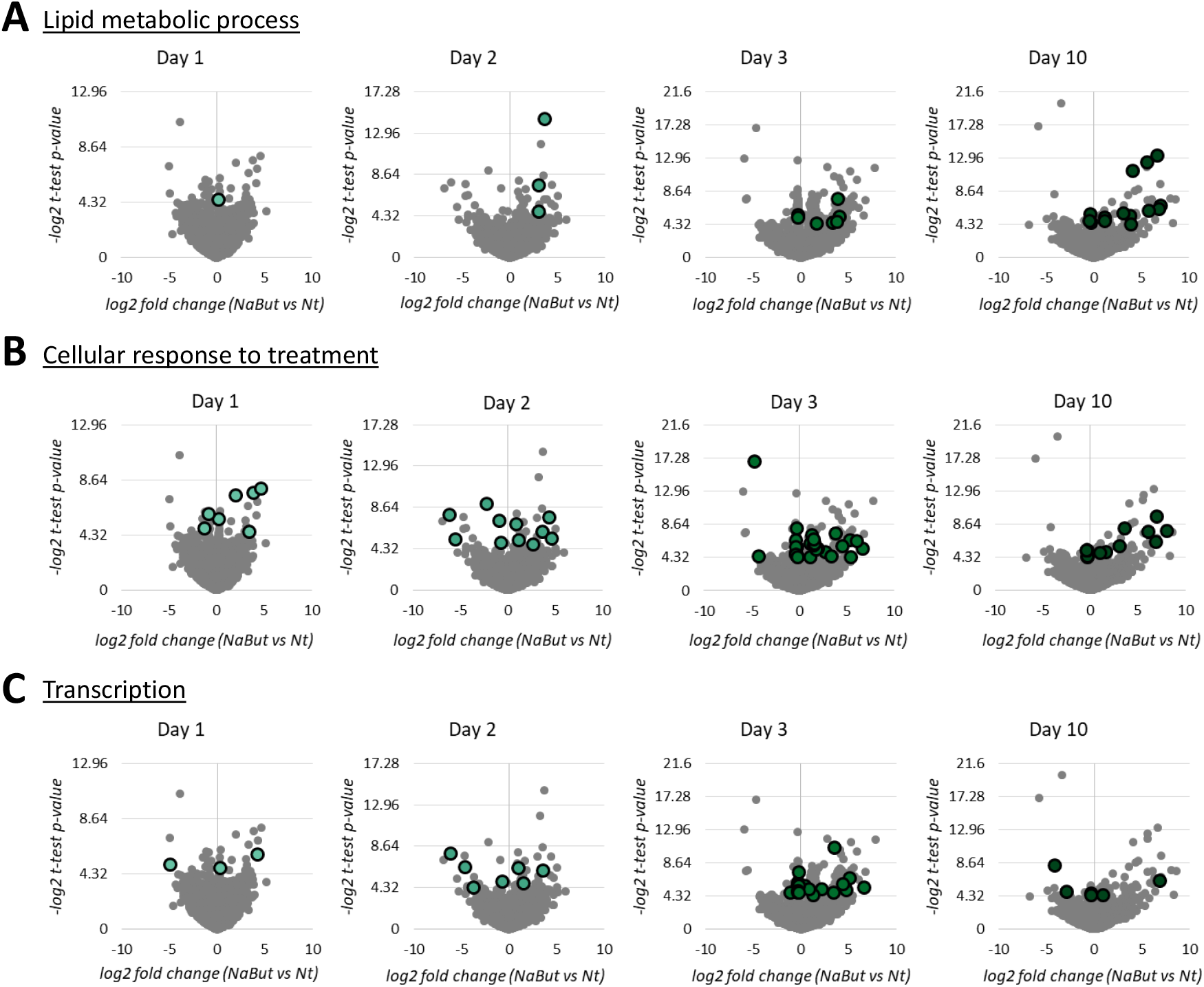
NaBut treatment affects the proteome of spheroids. Volcano plots representing NaBut vs Nt fold change after treatment (day 1, 2 and 3) and recovery period (day 10). Highlighted bubbles display the relative abundance of proteins related to **(A)** lipid metabolic process, **(B)** cellular response to treatment, and **(C)** transcription.

**Fig. 6.**
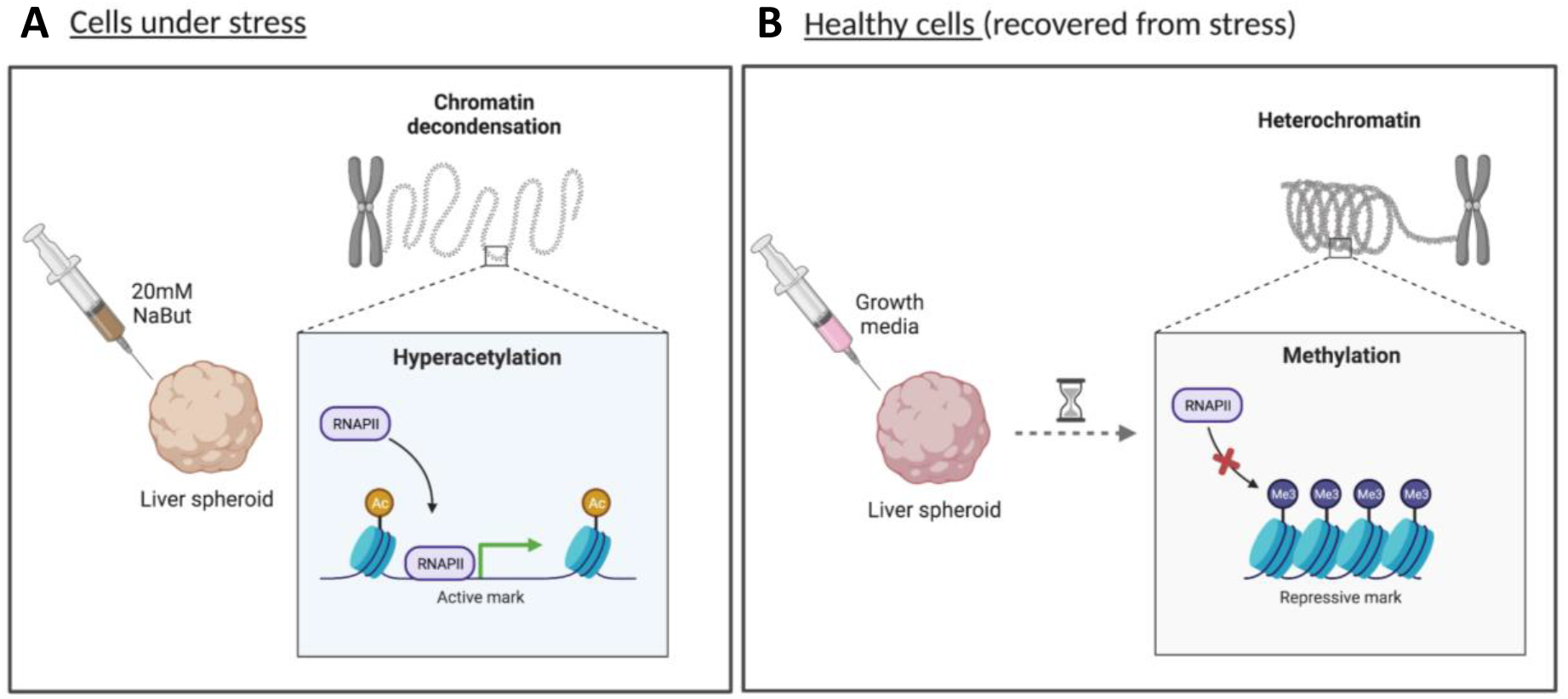
Schematic representation of direct and indirect targets of NaBut treatment on spheroids. **(A)** NaBut treatment induce direct effects such as histone hyperacetylation and indirect, such as chromatin decondensation and transcription activation. **(B)** Removal of treatment and reestablishment of standard culture condition revert the NaBut effects and cells return their normal physiology.

## Discussion

In the present study, we successfully established long-term liver spheroids as a functional system that closely resemble the phenotype of a solid liver tissue. We have used a cell culture system in which spheroids were cultured in incubators using the clinostat principle, i.e. rotating bioreactors. This approach provides an optimal growth environment allowing spheroids to be exposed to an equal and very low shear force while actively homogenizing nutrients in the volume of the bioreactor, resulting in a uniform culture [11]. In contrast to liver cell lines grown in 2D, spheroids growing in this dynamic system attach to one another and form 3D structures, resembling a parental tissue phenotype.

Given the relative novelty of this cell culture method, numerous applications still need to be explored. We aimed to demonstrate that cells grown as spheroids can be treated with epigenetics drugs at high concentration, i.e. 20 mM sodium butyrate, and still maintain their viability to investigate the response to the treatment and the recovery. Notably, the same treatment on 2D cells leads to rapid cell death in just a few days. Moreover, the slow proliferation of cells in 3D spheroids allows to investigate the recovery effect directly on the same cells that experienced the treatment, while with 2D culture 10 days is a sufficient time frame to produce a whole new layer of daughter cells.

Other aspects of 3D cell culture make this model convenient for our type of study. For instance, once in the bioreactor, spheroids are treated essentially the same way as cells maintained in 2D culture. However, one of the main advantages of this system is that, compared to flat culture, media exchange and cell harvesting is performed without the use of trypsinization maintaining unharmed all cell-cell communications and receptors. Although trypsinization is the most popular method used to detach cells from a culture surface, it decreases the number of functional integrins on the cell membrane and consequently reduces the cell ability to form adhesive bonds and communicate with each other [26]. HepG2/C3A spheroids used in our study were maintained for more than 30 days in culture to guarantee a metabolically competent and mature cell model. In fact, the liver spheroids described here were found to closely resemble liver tissues at the proteome level, which is in line with studies of human liver organoids, where high levels of liver-related proteins were identified [27–30]. We were able to detect in our 3D model an enrichment of important biomarkers as well as proteins from specific liver pathways that are critical for energy production [17, 31]. The liver is also the major organ for metal detoxification, of which function is mediated by metallothioneins (MT). MT-1 and -2 isoforms detected in this study play a protective role by binding heavy metals such as cadmium, zinc, and copper, preventing metal-induced oxidative damage [32, 33].

The reproduction of the parental tissue physiology and the long-term functionalities of our spheroids create great experimental tools for diverse research purposes, including drugs and chemicals [34, 35] screening. We tested the spheroids responsiveness to drug treatment and demonstrate that, upon exposure, they respond to stimulation by changing the metabolism and chromatin dynamics. By treating spheroids with a drug that yields acetyl-CoA metabolites, we observed changes in histone acetylation, which triggers chromatin state rearrangements [36]. The modulation of transcription through core histone acetylation is one of the most relevant mechanisms by which cell are epigenetically regulated [37]. On the other hand, we demonstrated that basal levels of histone acetylation can be restored in a few days, suggesting that this histone mark seems to have minimal epigenetics memory in the cell culture, as it is not being conserved once treatment has been removed.

As a final note, our 3D system is also relatively environmentally friendly, as a few spheroids are sufficient to perform a comprehensive proteomics and transcriptomics analysis. Therefore, a single 10 mL bioreactor can contain all the replicates and time points required for a complete experiment, while a 2D cell experiment would require a different petri dish for each replicate and time point. We did not demonstrate this aspect, but we speculate that this can also contribute to improving the reproducibility of experiments and maintain low costs of operation.

## Materials and Methods

### Flat cell culture

HepG2/C3A cell line (from human hepatocellular carcinoma) was obtained from the American Type Culture Collection (ATCC, CRL-10741). The cells were maintained in Dulbecco’s Modified Eagle’s Medium (DMEM, containing 4.5g/L glucose) supplemented with 10% Fetal Bovine Serum (FBS), 1% Non-Essential Amino Acids, 1% GlutaMAX and 0.5% Penicillin/Streptomycin (all Corning). Cells were kept in a controlled atmosphere (5% CO_2_ incubator at 37°C). Before use for spheroid cultures, flat cells were cultured until reach 80% confluence and were trypsinized using 0.5% Trypsin/EDTA diluted 1:1 in Hanks solution (Corning) for 5 minutes, followed by centrifugation at 140 g for 5 minutes. The number of cells was estimated using the Corning Cell Counter and CytoSMART Cloud App (Corning). Flat cells images were acquired with PAULA Smart Cell Imager (Leica) using a 10x magnification.

### Spheroid culture

Spheroids were prepared and cultured according to [10], with some modifications.

#### Preparation of spheroids

HepG2/C3A spheroids were prepared using an ultra-low attachment 24-well round bottom plate containing microwells (Elplasia®, Corning). Before use, wells were washed with growth media. Wells were then prefilled with growth media and the plate was centrifuged (3,000g for 5 min) to remove air bubbles from the well surface. Cell suspension (1.2×10^6^ cells diluted in growth media) was added to each Elplasia® well, followed by plate centrifugation (120 g for 3 min) and overnight incubation under a controlled atmosphere to allow cell self-aggregation and spheroids formation.

#### Spheroids culture into ClinoReactors

prior to use for spheroid culture, the water beads, inside the humidity chamber of the ClinoReactor (CelVivo), were hydrated with sterile water (Corning) and the growth chamber was filled with the proper growth media. The ClinoReactor was incubated for 24 h, rotating in the ClinoStar (CelVivo, Denmark), to equilibrate under controlled atmosphere (5% CO_2_ incubator at 37°C). Spheroids were detached from the Elplasia® well by gently pipetting up and down, followed by washing with pre-warmed growth media. The detached spheroids were collected into a Petri dish and their quality (compactness and roundness) checked under the microscope. Good quality spheroids were selected and transferred into equilibrated ClinoReactors filled with fresh growth media (day 1). ClinoReactors were placed in the ClinoStar and the rotation speed was set between 10-11 rpm. On day 15 the spheroids population growing in the ClinoReactor was split into two ClinoReactors to adjust the population density. Optimal growth conditions were achieved by exchanging media three times a week and adjusting rotation speed according to spheroids growth. Spheroids images were acquired with PAULA Smart Cell Imager (Leica) using a 10x magnification.

### Cell viability assay

Cell viability was assessed by measuring adenylate kinase (AK) levels using ToxiLight™ assay kit (Lonza). Briefly, HepG2/C3A flat cells and spheroids supernatants were collected in duplicate and 20 µL were transferred to a 96 well white-walled plate (flat bottom clear). For each assay plate, a standard curve was prepared using different amounts of cells (156 to 10,000 HepG2/C3A cells per well) lysed in digitonin lysis buffer (Promega). One hundred microliters of adenylate kinase detection reagent (Lonza) were added to each well and the content was homogenized by gently pipetting up and down. Bubbles in the assay were removed by centrifugation with pressure. The plate was incubated for 20 min at room temperature and read in a Victor X5 plate reader (Perkin Elmer), in luminescent mode. Results are shown as the percentage of dead cells, calculated using the assay reading divided by the number of cells per spheroid estimated by Wrzesinski and Fey [10].

### Metabolic labeling

For labeling experiment, HepG2/C3A flat cells (1×10^5^ cells/well, in a 96-well plate) and spheroids (28 days old) were used. Control spheroids (unlabeled) were collected before adding the heavy labeled media. Isotope-labeled arginine (^13^C_6_ ^15^N_4_, Sigma) was added to the standard growth media to 420mg/L and cells were cultured for 4 days. At the time points, cells were collected, washed with HBSS, and stored the proteins were extracted for mass spectrometry analysis.

### NaBut treatment

To evaluate the effect of a histone deacetylase inhibitor (HDACi) on the proteome and histone post-translational modifications, HepG2/C3A spheroids and flat cells were treated with 20 mM of sodium butyrate (NaBut). Spheroids were maintained in growth media, as described above, for 35 days. Prior to the assay, spheroids were divided into two ClinoReactors, corresponding to treatment and control groups. Flat cells were trypsinized and plated in a 96 well plate (2×10^5^ cells/well). After 24 h, growth media was replaced by growth media containing 20 mM of NaBut for the treatment group while the non-treated group received growth media only. Spheroids and flat cells were treated for 3 days and, after collection, the treatment media was replaced by the standard growth media and spheroids were maintained for additional 7 days, until the last collection on day 10. Spheroids were collected daily, washed with Hanks solution to remove the FBS and centrifuged at 1000 rpm for 5 min. The supernatant was discarded, and the dry pellets were immediately stored in −80°C until processing.

### Metabolite profile analysis

The supernatant of flat cells and spheroids treated with NaBut were collected and centrifuged at 1000 g for 5 minutes. The supernatant was transferred to a new tube and stored at −20°C until use. Twenty microliters of each supernatant were then mixed with four volumes of methanol containing internal standards. After vortexing, samples were centrifuged at 14000 rpm for 10 minutes and the supernatant was transferred into glass vials. Samples were analyzed with ABsciex 6500+ with Ace PFP column. A pooled quality control (QC) sample was also added to the sample list. The QC sample was injected six times for coefficient of variation (CV) calculation and data quality control.

### Protein extraction and sample preparation

To analyze the proteome, cytoplasm and nuclei of both flat cells and spheroids pellets were isolated. Briefly, the pellets were resuspended with cold buffer A (10 mM ammonium bicarbonate pH 8, 1.5 mM MgCl_2_, and 10 mM KCl), centrifuged for 5 min at 400 g and the supernatant was removed. The pellet was resuspended in buffer A containing 10 mM of sodium butyrate, 0.15% NP-40 (v/v) and protease inhibitors. The suspension was centrifuged for 15 min at 3,200 g at 4°C and the supernatant (cytoplasm) was transferred to a new tube. The pellet was washed with PBS, centrifuged for 5 min at 3,200g and the supernatant was discarded. The final pellet consisted of crude nuclei. Proteins from the cytoplasm and nuclear fraction were digested using S-Trap filters (Protifi), according to the manufacturer’s instructions. Briefly, the samples were mixed with 5% SDS, followed by incubation with 5 mM DTT for 1 h and 20 mM iodoacetamide for 30 min in the dark, in order to reduce and alkylate proteins. Afterwards, phosphoric acid was added to the samples at a final concentration of 1.2%. Samples were diluted in six volumes of binding buffer (90 % methanol and 10 mM ammonium bicarbonate, pH 8.0). After gentle mixing, the protein solution was loaded to an S-Trap filter and spun at 500 g for 30 sec. The samples were washed twice with binding buffer. Finally, 1 µg of sequencing grade trypsin (Promega), diluted in 50 Mm ammonium bicarbonate, was added into the S-trap filter and samples were digested overnight at 37°C. Peptides were eluted in three steps: (i) 40 µl of 50 mM ammonium bicarbonate, (ii) 40 µl of 0.1% trifluoroacetic acid (TFA) and (iii) 40 µl of 60% acetonitrile and 0.1% TFA. The peptide solution was pooled, spun at 1,000 g for 30 sec and dried in a vacuum centrifuge.

### Histone extraction and sample preparation

Histone proteins were extracted from the cell pellet as described by [38] to ensure good-quality identification and quantification of single histone marks, what would not be possible by extracting these data from proteomics runs. Briefly, histones were acid-extracted with chilled 0.2M sulfuric acid (5:1, sulfuric acid : pellet) and incubated with constant rotation for 4 h at 4°C, followed by precipitation with 33% trichloroacetic acid (TCA) overnight at 4°C. Then, the supernatant was removed, and the tubes were rinsed with ice-cold acetone containing 0.1% HCl, centrifuged and rinsed again using 100% ice-cold acetone. After the final centrifugation, the supernatant was discarded, and the pellet was dried using a vacuum centrifuge. The pellet was dissolved in 50 mM ammonium bicarbonate, pH 8.0. In the fume hood, samples were mixed with 5 µL of acetonitrile, followed by 5 µL of propionic anhydride and 14 µL of ammonium hydroxide (all Sigma Aldrich) to balance the pH at 8.0. The mixture was incubated for 15 min and the procedure was repeated. Histones were then digested with 1 µg of sequencing grade trypsin (Promega) diluted in 50 mM ammonium bicarbonate (1:20, enzyme:sample) overnight at room temperature. Derivatization reaction was repeated to derivatize peptide N-termini. The samples were dried in a vacuum centrifuge.

### Sample desalting

Prior to mass spectrometry analysis, samples were desalted using a 96-well plate filter (Orochem) packed with 1 mg of Oasis HLB C-18 resin (Waters). Briefly, the samples were resuspended in 100 µl of 0.1% TFA and loaded onto the HLB resin, which was previously equilibrated using 100 µl of the same buffer. After washing with 100 µl of 0.1% TFA, the samples were eluted with a buffer containing 70 µl of 60% acetonitrile and 0.1% TFA and then dried in a vacuum centrifuge.

### LC-MS/MS Acquisition

Samples were resuspended in 10 µl of 0.1% TFA and loaded onto a Dionex RSLC Ultimate 300 (Thermo Scientific), coupled online with an Orbitrap Fusion Lumos (Thermo Scientific). Chromatographic separation was performed with a two-column system, consisting of a C-18 trap cartridge (300 µm ID, 5 mm length) and a picofrit analytical column (75 µm ID, 25 cm length) packed in-house with reversed-phase Repro-Sil Pur C18-AQ 3 µm resin. To analyze the proteome, peptides were separated using a 180 min gradient from 4-30% buffer B (buffer A: 0.1% formic acid, buffer B: 80% acetonitrile + 0.1% formic acid) at a flow rate of 300nl/min. The mass spectrometer was set to acquire spectra in a data-dependent acquisition (DDA) mode. Briefly, the full MS scan was set to 300-1200 m/z in the orbitrap with a resolution of 120,000 (at 200 m/z) and an AGC target of 5×10e5. MS/MS was performed in the ion trap using the top speed mode (2 sec), and AGC target of 1×10e4 and an HCD collision energy of 35.

To analyze the histones, peptides were separated using a 60 min gradient from 4-30% buffer B (buffer A: 0.1% formic acid, buffer B: 80% acetonitrile + 0.1% formic acid) at a flow rate of 300nl/min. The mass spectrometer was set to acquire spectra in a data-independent acquisition (DIA) mode. Briefly, the full MS scan was set to 300-1100 m/z in the orbitrap with a resolution of 120,000 (at 200 m/z) and an AGC target of 5×10e5. MS/MS was performed in the orbitrap with sequential isolation windows of 50 m/z with an AGC target of 2×10e5 and an HCD collision energy of 30.

### Proteomics and histone data analysis

Proteome raw files were searched using Proteome Discoverer software (v2.4, Thermo Scientific) using SEQUEST search engine and the SwissProt human database (updated February 2020). The search for total proteome included variable modification of N-terminal acetylation and fixed modification of carbamidomethyl cysteine. Trypsin was specified as the digestive enzyme with two missed cleavages allowed. Mass tolerance was set to 10 ppm for precursor ions and 0.2 Da for product ions. Peptide and protein false discovery rate was set to 1%. Each analysis was performed with two biological replicates. Prior statistics, proteins were log2 transformed, normalized by the average value of each sample and missing values were imputed using a normal distribution 2 standard deviations lower than the mean as described [39]. Although the protein extraction was done by isolating cytoplasm and nuclei, they were used as replicates and an average was calculated for the analysis. Statistical regulation was assessed using heteroscedastic T-test (if p-value < 0.05). Data distribution was assumed to be normal, but this was not formally tested.

Histone peptides raw files were imported into EpiProfile 2.0 software [22]. From the extracted ion chromatogram, the area under the curve was obtained and used to estimate the abundance of each peptide. In order to achieve the relative abundance of post-translational modifications (PTMs), the sum of all different modified forms of a histone peptide was considered as 100% and the area of the particular peptide was divided by the total area for that histone peptide in all of its modified forms. The relative ratio of two isobaric forms was estimated by averaging the ratio for each fragment ion with different mass between the two species. The resulting peptide lists generated by EpiProfile were exported to Microsoft Excel and further processed for a detailed analysis.

### RNA extraction and sequencing

5 spheroids (how many days old) stored at −80C were thawed on ice and then mechanically homogenized. RNA was extracted using an RNeasy micro kit (Qiagen). RNA purity and concentration was measured using a nanodrop and samples with a 260/280 ratio greater than 2 were kept. RNA integrity number (RIN) was then measured using a Bioanalyzer and samples with a RIN greater than 8 were kept. RNA was then sent to Novogene for library preparation and whole transcriptome sequencing. Libraries were sequenced on the NovaSeq platform to obtain an average of ∼45 million paired-end 150 bp reads per sample.

### RNA-seq data analysis

Raw reads were trimmed to remove low quality base calls and Illumina universal adapters using Trim Galore! (Version 0.6.5) with default parameters and then assessed using fastQC (version 0.11.4) and multiqc (version 1.10.1). Reads were then aligned to the human genome (GRCh38) using STAR with default parameters. Alignment quality control was performed using RSeQC and Qualimap. Quantification was performed using RSEM. Quantification quality control was performed using EDASeq (version 2.3) and NOISeq (version 2.4). Time-course differential expression analysis was performed using msSigPro (version 1.68). Clustering of differential time-course genes was performed by identifying the optimal number of clusters using mclust (version 5.4.1) and then clustering using k-means method.

Gene ontology analysis was performed using clusterProfiler (version 4.4.4) where cell type enrichments utilized MSigDB (version 7.5.1). The code used for the analysis is provided as a supplementary file.

### Mass spectrometry raw data availability

All raw mass spectrometry data files from this study have been submitted to the Chorus repository (https://chorusproject.org/pages/index.html) under project number 1786.

### RNA-seq raw data availability

All raw and processed RNA-Seq data files from this study have been submitted to the gene expression omnibus repository under accession number GSE213944.

## Supporting information

Code for RNA-seq analysis

## Acknowledgments

The Sidoli lab would like to acknowledge the team of CelVivo IVS (Odense, Denmark) for all the support with establishing the 3D spheroids culture, in particular Dr. Krzysztof Wrzesinski, Dr. Peter Willems-Alnøe and Dr. Stephen J. Fey. A thank you goes also to Ms. Georgia Fallon (now Dr. Jonathan Lai’s lab, Einstein) for the initial contribution with the introduction of the manuscript. We also thank the staff from the Stable Isotope and Metabolomics Core Facility of the Diabetes Research and Training Center (DRTC) of the Albert Einstein College of Medicine.

## Authors’ contributions

Study conception and design: S.Str., E.N., S.Sid.. Experiments and procedures: S.Str., R.C., J.A.. Analysis and interpretation of data: S.Str., R.C., S.Sid.. Drafting of the manuscript: S.Str., S.Sid.. All authors reviewed the manuscript.

## References

1. Antoni, D., et al., Three-dimensional cell culture: a breakthrough in vivo. Int J Mol Sci, 2015. 16(3): p. 5517–27.

2. Kapałczyńska, M., et al., 2D and 3D cell cultures - a comparison of different types of cancer cell cultures. Arch Med Sci, 2018. 14(4): p. 910–919.

3. Lee, J.M., et al., A three-dimensional microenvironment alters protein expression and chemosensitivity of epithelial ovarian cancer cells in vitro. Lab Invest, 2013. 93(5): p. 528–42.

4. Wu, Y.M., et al., Morphological changes and molecular expressions of hepatocellular carcinoma cells in three-dimensional culture model. Exp Mol Pathol, 2009. 87(2): p. 133–40.

5. Kim, J.B., Three-dimensional tissue culture models in cancer biology. Semin Cancer Biol, 2005. 15(5): p. 365–77.

6. Wrzesinski, K., et al., The cultural divide: exponential growth in classical 2D and metabolic equilibrium in 3D environments. PLoS One, 2014. 9(9): p. e106973.

7. Edmondson, R., et al., Three-dimensional cell culture systems and their applications in drug discovery and cell-based biosensors. Assay Drug Dev Technol, 2014. 12(4): p. 207–18.

8. Gonda, S.R., et al., Three-dimensional transgenic cell model to quantify genotoxic effects of space environment. Adv Space Res, 2001. 27(2): p. 421–30.

9. Wrzesinski, K., et al., HepG2/C3A 3D spheroids exhibit stable physiological functionality for at least 24 days after recovering from trypsinisation. 2013, Toxicology Research. p. 163–172.

10. Wrzesinski, K. and S.J. Fey, After trypsinisation, 3D spheroids of C3A hepatocytes need 18 days to re-establish similar levels of key physiological functions to those seen in the liver. 2013, Toxicology Research. p. 123–135.

11. Wrzesinski, K. and S.J. Fey, Metabolic Reprogramming and the Recovery of Physiological Functionality in 3D Cultures in Micro-Bioreactors. Bioengineering (Basel), 2018. 5(1).

12. Tvardovskiy, A., et al., Top-down and Middle-down Protein Analysis Reveals that Intact and Clipped Human Histones Differ in Post-translational Modification Patterns. Mol Cell Proteomics, 2015. 14(12): p. 3142–53.

13. Sidoli, S., et al., A mass spectrometry-based assay using metabolic labeling to rapidly monitor chromatin accessibility of modified histone proteins. Sci Rep, 2019. 9(1): p. 13613.

14. Feist, P.E., et al., Multicellular Tumor Spheroids Combined with Mass Spectrometric Histone Analysis To Evaluate Epigenetic Drugs. Anal Chem, 2017. 89(5): p. 2773–2781.

15. Li, J., et al., Hi-C profiling of cancer spheroids identifies 3D-growth-specific chromatin interactions in breast cancer endocrine resistance. Clin Epigenetics, 2021. 13(1): p. 175.

16. Babula, P., et al., Mammalian metallothioneins: properties and functions. Metallomics, 2012. 4(8): p. 739–50.

17. Chiang, J., Liver Physiology: Metabolism and Detoxification, in Pathobiology of Human Disease, R.N.M. Linda M. McManus, Editor. 2014, Elsevier: San Diego. p. 1770–1782.

18. Shannon, P., et al., Cytoscape: a software environment for integrated models of biomolecular interaction networks. Genome Res, 2003. 13(11): p. 2498–504.

19. Candido, E.P., R. Reeves, and J.R. Davie, Sodium butyrate inhibits histone deacetylation in cultured cells. Cell, 1978. 14(1): p. 105–13.

20. Eckschlager, T., et al., Histone Deacetylase Inhibitors as Anticancer Drugs. Int J Mol Sci, 2017. 18(7).

21. Sidoli, S. and B.A. Garcia, Characterization of Individual Histone Posttranslational Modifications and Their Combinatorial Patterns by Mass Spectrometry-Based Proteomics Strategies. Methods Mol Biol, 2017. 1528: p. 121–148.

22. Yuan, Z.F., et al., EpiProfile 2.0: A Computational Platform for Processing EpiProteomics Mass Spectrometry Data. J Proteome Res, 2018. 17(7): p. 2533–2541.

23. Suarez-Alvarez, B., et al., DNA methylation: a promising landscape for immune system-related diseases. Trends Genet, 2012. 28(10): p. 506–14.

24. Bannister, A.J. and T. Kouzarides, Regulation of chromatin by histone modifications. Cell Res, 2011. 21(3): p. 381–95.

25. Eden, E., et al., GOrilla: a tool for discovery and visualization of enriched GO terms in ranked gene lists. BMC Bioinformatics, 2009. 10: p. 48.

26. Brown, M.A., et al., The use of mild trypsinization conditions in the detachment of endothelial cells to promote subsequent endothelialization on synthetic surfaces. Biomaterials, 2007. 28(27): p. 3928–35.

27. Mun, S.J., et al., Generation of expandable human pluripotent stem cell-derived hepatocyte-like liver organoids. J Hepatol, 2019. 71(5): p. 970–985.

28. Bell, C.C., et al., Characterization of primary human hepatocyte spheroids as a model system for drug-induced liver injury, liver function and disease. Sci Rep, 2016. 6: p. 25187.

29. Broutier, L., et al., Human primary liver cancer-derived organoid cultures for disease modeling and drug screening. Nat Med, 2017. 23(12): p. 1424–1435.

30. Sun, L., et al., Modelling liver cancer initiation with organoids derived from directly reprogrammed human hepatocytes. Nat Cell Biol, 2019. 21(8): p. 1015–1026.

31. Begriche, K., et al., Drug-induced toxicity on mitochondria and lipid metabolism: mechanistic diversity and deleterious consequences for the liver. J Hepatol, 2011. 54(4): p. 773–94.

32. Davis, S.R. and R.J. Cousins, Metallothionein expression in animals: a physiological perspective on function. J Nutr, 2000. 130(5): p. 1085–8.

33. Vasák, M., Advances in metallothionein structure and functions. J Trace Elem Med Biol, 2005. 19(1): p. 13–7.

34. Štampar, M., et al., Hepatocellular carcinoma (HepG2/C3A) cell-based 3D model for genotoxicity testing of chemicals. Sci Total Environ, 2021. 755(Pt 2): p. 143255.

35. Fey, S.J. and K. Wrzesinski, Determination of drug toxicity using 3D spheroids constructed from an immortal human hepatocyte cell line. Toxicol Sci, 2012. 127(2): p. 403–11.

36. Thomas, S.P. and J.M. Denu, Short-chain fatty acids activate acetyltransferase p300. Elife, 2021. 10.

37. Biancotto, C., G. Frigè, and S. Minucci, Histone modification therapy of cancer. Adv Genet, 2010. 70: p. 341–86.

38. Sidoli, S., et al., Complete Workflow for Analysis of Histone Post-translational Modifications Using Bottom-up Mass Spectrometry: From Histone Extraction to Data Analysis. J Vis Exp, 2016(111).

39. Aguilan, J.T., K. Kulej, and S. Sidoli, Guide for protein fold change and p-value calculation for non-experts in proteomics. Mol Omics, 2020. 16(6): p. 573–582.

