## Supplementary Material for "Investigation of reversible histone acetylation and dynamics in gene expression regulation using 3D liver spheroid model"

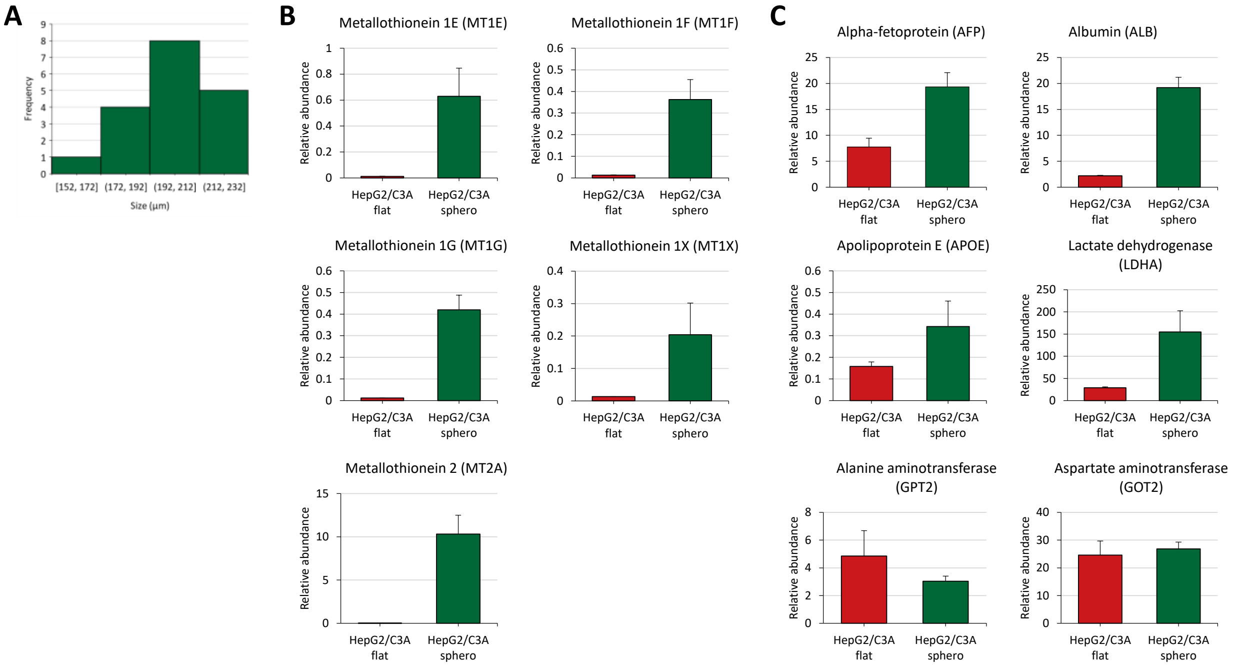


**Fig. S1. Spheroids express proteins characteristic of the human liver. (A)** Size distribution of liver spheroids. For the proteome analysis, flat cells and spheroids were collected, processed and the peptides analyzed by mass spectrometry. Bar graphs show the relative abundance of **(B)** metallothioneins and **(C)** liver biomarkers. Data are represented as means ± SEM.

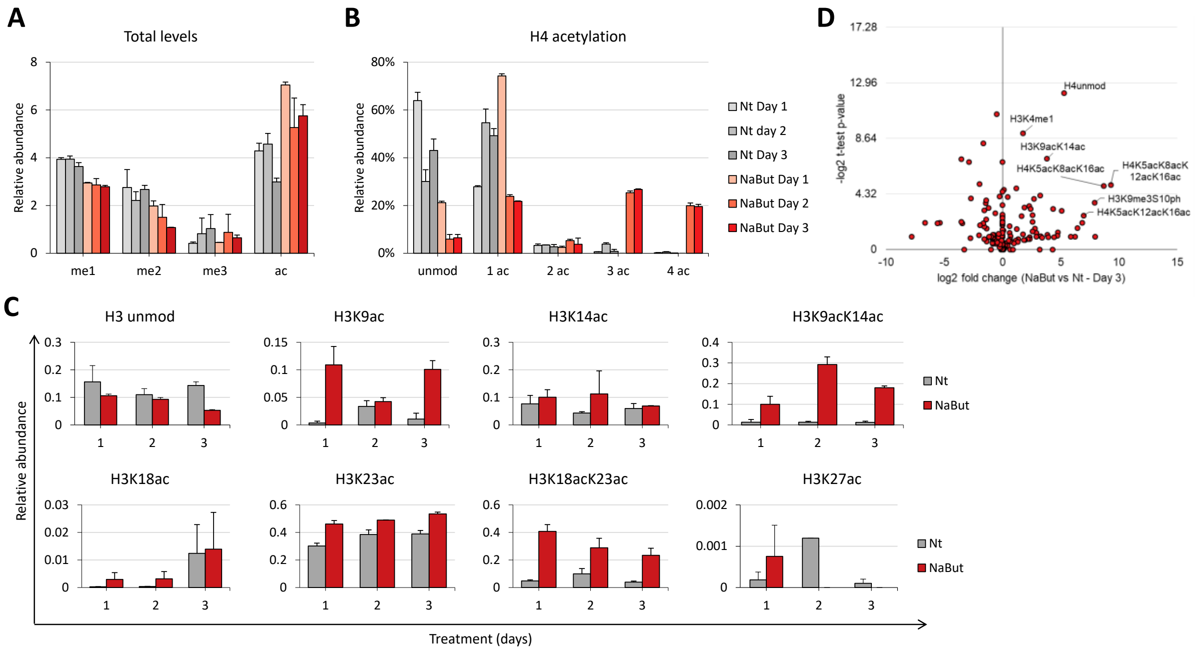


**Fig. S2. NaBut treatment induce histone hyperacetylation in flat cells.** HepG2/C3A cells were treated with 20 mM NaBut and were kept in culture for 3 days. After treatment, histones were extracted and analyzed by mass spectrometry. **(A)** Total levels of histone peptides containing 1, 2 or 3 methylations (me1, me2, me3, respectively) or containing acetylations (ac). **(B)** Total levels of histone H4 peptides containing acetylations (1ac, 2ac, 3ac, 4ac). Unmod, unmodified peptide. **(C)** Relative abundance of histone H3 acetylated peptides. Data are represented as means ± SD. Nt, non-treated. **(D)** Volcano plot representing NaBut vs Nt fold change after 3 days of treatment.

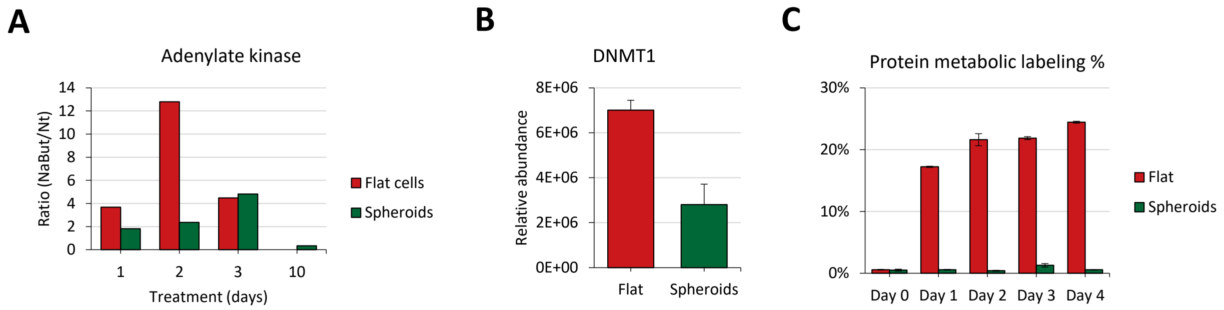


**Fig. S3. Liver spheroids have a slow replication rate and can recover from treatment. (A)** Adenylate kinase was measured following NaBut treatment. The culture supernatant from flat cells and spheroids were collected and analyzed by luminescence. **(B)** Relative abundance of DNMT1 in flat cells and spheroids. **(C)** Labeling incorporation in flat cells and spheroids. Data are represented as means ± SEM.

**Supplementary Table 1. Full metabolite profiling of the cell culture supernatant.**

**Supplementary Table 2. Gene expression clustering.**

**Supplementary Table 3. Proteome of HepG2/C3A spheroids and flat cells.**
