## Supplementary material for "Investigation of reversible histone acetylation and dynamics in gene expression regulation using 3D liver spheroid model": Code for RNA-seq analysis: Supplementary file_Code for RNA-Seq analysis.html

Protein Coding Time-course analysis


### Protein Coding Time-course analysis

Load Libraries

```
library(DESeq2)
```

```
## Loading required package: S4Vectors
```

```
## Warning: package 'S4Vectors' was built under R version 4.1.3
```

```
## Loading required package: stats4
```

```
## Loading required package: BiocGenerics
```

```
## 
## Attaching package: 'BiocGenerics'
```

```
## The following objects are masked from 'package:stats':
## 
##     IQR, mad, sd, var, xtabs
```

```
## The following objects are masked from 'package:base':
## 
##     anyDuplicated, append, as.data.frame, basename, cbind, colnames,
##     dirname, do.call, duplicated, eval, evalq, Filter, Find, get, grep,
##     grepl, intersect, is.unsorted, lapply, Map, mapply, match, mget,
##     order, paste, pmax, pmax.int, pmin, pmin.int, Position, rank,
##     rbind, Reduce, rownames, sapply, setdiff, sort, table, tapply,
##     union, unique, unsplit, which.max, which.min
```

```
## 
## Attaching package: 'S4Vectors'
```

```
## The following objects are masked from 'package:base':
## 
##     expand.grid, I, unname
```

```
## Loading required package: IRanges
```

```
## Loading required package: GenomicRanges
```

```
## Loading required package: GenomeInfoDb
```

```
## Loading required package: SummarizedExperiment
```

```
## Loading required package: MatrixGenerics
```

```
## Loading required package: matrixStats
```

```
## 
## Attaching package: 'MatrixGenerics'
```

```
## The following objects are masked from 'package:matrixStats':
## 
##     colAlls, colAnyNAs, colAnys, colAvgsPerRowSet, colCollapse,
##     colCounts, colCummaxs, colCummins, colCumprods, colCumsums,
##     colDiffs, colIQRDiffs, colIQRs, colLogSumExps, colMadDiffs,
##     colMads, colMaxs, colMeans2, colMedians, colMins, colOrderStats,
##     colProds, colQuantiles, colRanges, colRanks, colSdDiffs, colSds,
##     colSums2, colTabulates, colVarDiffs, colVars, colWeightedMads,
##     colWeightedMeans, colWeightedMedians, colWeightedSds,
##     colWeightedVars, rowAlls, rowAnyNAs, rowAnys, rowAvgsPerColSet,
##     rowCollapse, rowCounts, rowCummaxs, rowCummins, rowCumprods,
##     rowCumsums, rowDiffs, rowIQRDiffs, rowIQRs, rowLogSumExps,
##     rowMadDiffs, rowMads, rowMaxs, rowMeans2, rowMedians, rowMins,
##     rowOrderStats, rowProds, rowQuantiles, rowRanges, rowRanks,
##     rowSdDiffs, rowSds, rowSums2, rowTabulates, rowVarDiffs, rowVars,
##     rowWeightedMads, rowWeightedMeans, rowWeightedMedians,
##     rowWeightedSds, rowWeightedVars
```

```
## Loading required package: Biobase
```

```
## Welcome to Bioconductor
## 
##     Vignettes contain introductory material; view with
##     'browseVignettes()'. To cite Bioconductor, see
##     'citation("Biobase")', and for packages 'citation("pkgname")'.
```

```
## 
## Attaching package: 'Biobase'
```

```
## The following object is masked from 'package:MatrixGenerics':
## 
##     rowMedians
```

```
## The following objects are masked from 'package:matrixStats':
## 
##     anyMissing, rowMedians
```

```
library(tximport)
library(affy)
library(ggplot2)
library(vsn)
library(ggrepel)
library(factoextra)
```

```
## Welcome! Want to learn more? See two factoextra-related books at https://goo.gl/ve3WBa
```

```
library(FactoMineR)
library(genefilter)
```

```
## 
## Attaching package: 'genefilter'
```

```
## The following objects are masked from 'package:MatrixGenerics':
## 
##     rowSds, rowVars
```

```
## The following objects are masked from 'package:matrixStats':
## 
##     rowSds, rowVars
```

```
#library(fdrtool)
library(reshape2)
library(RColorBrewer)
library(biomaRt)
library(EnhancedVolcano)
```

```
## Registered S3 methods overwritten by 'ggalt':
##   method                  from   
##   grid.draw.absoluteGrob  ggplot2
##   grobHeight.absoluteGrob ggplot2
##   grobWidth.absoluteGrob  ggplot2
##   grobX.absoluteGrob      ggplot2
##   grobY.absoluteGrob      ggplot2
```

```
library(readxl)
library(pheatmap)
library(zFPKM)
library(umap)
library(ComplexHeatmap)
```

```
## Loading required package: grid
```

```
## ========================================
## ComplexHeatmap version 2.10.0
## Bioconductor page: http://bioconductor.org/packages/ComplexHeatmap/
## Github page: https://github.com/jokergoo/ComplexHeatmap
## Documentation: http://jokergoo.github.io/ComplexHeatmap-reference
## 
## If you use it in published research, please cite:
## Gu, Z. Complex heatmaps reveal patterns and correlations in multidimensional 
##   genomic data. Bioinformatics 2016.
## 
## The new InteractiveComplexHeatmap package can directly export static 
## complex heatmaps into an interactive Shiny app with zero effort. Have a try!
## 
## This message can be suppressed by:
##   suppressPackageStartupMessages(library(ComplexHeatmap))
## ========================================
## ! pheatmap() has been masked by ComplexHeatmap::pheatmap(). Most of the arguments
##    in the original pheatmap() are identically supported in the new function. You 
##    can still use the original function by explicitly calling pheatmap::pheatmap().
```

```
## 
## Attaching package: 'ComplexHeatmap'
```

```
## The following object is masked from 'package:pheatmap':
## 
##     pheatmap
```

```
## The following object is masked from 'package:genefilter':
## 
##     dist2
```

```
library(sva)
```

```
## Loading required package: mgcv
```

```
## Loading required package: nlme
```

```
## 
## Attaching package: 'nlme'
```

```
## The following object is masked from 'package:IRanges':
## 
##     collapse
```

```
## This is mgcv 1.8-40. For overview type 'help("mgcv-package")'.
```

```
## Loading required package: BiocParallel
```

```
library(data.table)
```

```
## 
## Attaching package: 'data.table'
```

```
## The following objects are masked from 'package:reshape2':
## 
##     dcast, melt
```

```
## The following object is masked from 'package:SummarizedExperiment':
## 
##     shift
```

```
## The following object is masked from 'package:GenomicRanges':
## 
##     shift
```

```
## The following object is masked from 'package:IRanges':
## 
##     shift
```

```
## The following objects are masked from 'package:S4Vectors':
## 
##     first, second
```

```
library(purrr)
```

```
## 
## Attaching package: 'purrr'
```

```
## The following object is masked from 'package:data.table':
## 
##     transpose
```

```
## The following object is masked from 'package:GenomicRanges':
## 
##     reduce
```

```
## The following object is masked from 'package:IRanges':
## 
##     reduce
```

```
library(dplyr)
```

```
## 
## Attaching package: 'dplyr'
```

```
## The following objects are masked from 'package:data.table':
## 
##     between, first, last
```

```
## The following object is masked from 'package:nlme':
## 
##     collapse
```

```
## The following object is masked from 'package:biomaRt':
## 
##     select
```

```
## The following object is masked from 'package:Biobase':
## 
##     combine
```

```
## The following object is masked from 'package:matrixStats':
## 
##     count
```

```
## The following objects are masked from 'package:GenomicRanges':
## 
##     intersect, setdiff, union
```

```
## The following object is masked from 'package:GenomeInfoDb':
## 
##     intersect
```

```
## The following objects are masked from 'package:IRanges':
## 
##     collapse, desc, intersect, setdiff, slice, union
```

```
## The following objects are masked from 'package:S4Vectors':
## 
##     first, intersect, rename, setdiff, setequal, union
```

```
## The following objects are masked from 'package:BiocGenerics':
## 
##     combine, intersect, setdiff, union
```

```
## The following objects are masked from 'package:stats':
## 
##     filter, lag
```

```
## The following objects are masked from 'package:base':
## 
##     intersect, setdiff, setequal, union
```

```
library(tidyr)
```

```
## 
## Attaching package: 'tidyr'
```

```
## The following object is masked from 'package:reshape2':
## 
##     smiths
```

```
## The following object is masked from 'package:S4Vectors':
## 
##     expand
```

```
library(viridis)
```

```
## Loading required package: viridisLite
```

```
library(maSigPro)

colors <- viridis(2)
```

Load counts

```
dir <- "/Users/ronaldcutler/Library/CloudStorage//My\ Drive/Sidoli_Lab/Projects/Chromatin\ Decondensation/Data/RNA/Counts/Gencode"

sampleTable <- as.data.frame(read_excel("/Users/ronaldcutler/Library/CloudStorage//My\ Drive/Sidoli_Lab/Projects/Chromatin\ Decondensation/Analysis/sample_table.xlsx"))
sampleTable$time <- as.factor(sampleTable$time)
sampleTable$conditiontime <- as.factor(paste(sampleTable$condition, sampleTable$time, sep = "-"))

files <- file.path(dir, sampleTable$sample_ID)
files <- paste0(files, ".genes.results")
names(files) <- sampleTable$sample
files
```

```
##                                                                                                                                                                          Nt1_1 
##  "/Users/ronaldcutler/Library/CloudStorage//My Drive/Sidoli_Lab/Projects/Chromatin Decondensation/Data/RNA/Counts/Gencode/C1_1.genes.results" 
##                                                                                                                                                                          Nt1_2 
##  "/Users/ronaldcutler/Library/CloudStorage//My Drive/Sidoli_Lab/Projects/Chromatin Decondensation/Data/RNA/Counts/Gencode/C1_2.genes.results" 
##                                                                                                                                                                          Nt1_3 
##  "/Users/ronaldcutler/Library/CloudStorage//My Drive/Sidoli_Lab/Projects/Chromatin Decondensation/Data/RNA/Counts/Gencode/C1_3.genes.results" 
##                                                                                                                                                                          Nt2_1 
##  "/Users/ronaldcutler/Library/CloudStorage//My Drive/Sidoli_Lab/Projects/Chromatin Decondensation/Data/RNA/Counts/Gencode/C2_1.genes.results" 
##                                                                                                                                                                          Nt2_2 
##  "/Users/ronaldcutler/Library/CloudStorage//My Drive/Sidoli_Lab/Projects/Chromatin Decondensation/Data/RNA/Counts/Gencode/C2_2.genes.results" 
##                                                                                                                                                                          Nt2_3 
##  "/Users/ronaldcutler/Library/CloudStorage//My Drive/Sidoli_Lab/Projects/Chromatin Decondensation/Data/RNA/Counts/Gencode/C2_3.genes.results" 
##                                                                                                                                                                          Nt3_1 
##  "/Users/ronaldcutler/Library/CloudStorage//My Drive/Sidoli_Lab/Projects/Chromatin Decondensation/Data/RNA/Counts/Gencode/C3_1.genes.results" 
##                                                                                                                                                                          Nt3_2 
##  "/Users/ronaldcutler/Library/CloudStorage//My Drive/Sidoli_Lab/Projects/Chromatin Decondensation/Data/RNA/Counts/Gencode/C3_2.genes.results" 
##                                                                                                                                                                          Nt3_3 
##  "/Users/ronaldcutler/Library/CloudStorage//My Drive/Sidoli_Lab/Projects/Chromatin Decondensation/Data/RNA/Counts/Gencode/C3_3.genes.results" 
##                                                                                                                                                                         Nt10_1 
## "/Users/ronaldcutler/Library/CloudStorage//My Drive/Sidoli_Lab/Projects/Chromatin Decondensation/Data/RNA/Counts/Gencode/C10_1.genes.results" 
##                                                                                                                                                                         Nt10_2 
## "/Users/ronaldcutler/Library/CloudStorage//My Drive/Sidoli_Lab/Projects/Chromatin Decondensation/Data/RNA/Counts/Gencode/C10_2.genes.results" 
##                                                                                                                                                                         Nt10_3 
## "/Users/ronaldcutler/Library/CloudStorage//My Drive/Sidoli_Lab/Projects/Chromatin Decondensation/Data/RNA/Counts/Gencode/C10_3.genes.results" 
##                                                                                                                                                                       NaBut1_1 
##  "/Users/ronaldcutler/Library/CloudStorage//My Drive/Sidoli_Lab/Projects/Chromatin Decondensation/Data/RNA/Counts/Gencode/N1_1.genes.results" 
##                                                                                                                                                                       NaBut1_2 
##  "/Users/ronaldcutler/Library/CloudStorage//My Drive/Sidoli_Lab/Projects/Chromatin Decondensation/Data/RNA/Counts/Gencode/N1_2.genes.results" 
##                                                                                                                                                                       NaBut1_3 
##  "/Users/ronaldcutler/Library/CloudStorage//My Drive/Sidoli_Lab/Projects/Chromatin Decondensation/Data/RNA/Counts/Gencode/N1_3.genes.results" 
##                                                                                                                                                                       NaBut2_1 
##  "/Users/ronaldcutler/Library/CloudStorage//My Drive/Sidoli_Lab/Projects/Chromatin Decondensation/Data/RNA/Counts/Gencode/N2_1.genes.results" 
##                                                                                                                                                                       NaBut2_2 
##  "/Users/ronaldcutler/Library/CloudStorage//My Drive/Sidoli_Lab/Projects/Chromatin Decondensation/Data/RNA/Counts/Gencode/N2_2.genes.results" 
##                                                                                                                                                                       NaBut2_3 
##  "/Users/ronaldcutler/Library/CloudStorage//My Drive/Sidoli_Lab/Projects/Chromatin Decondensation/Data/RNA/Counts/Gencode/N2_3.genes.results" 
##                                                                                                                                                                       NaBut3_1 
##  "/Users/ronaldcutler/Library/CloudStorage//My Drive/Sidoli_Lab/Projects/Chromatin Decondensation/Data/RNA/Counts/Gencode/N3_1.genes.results" 
##                                                                                                                                                                       NaBut3_2 
##  "/Users/ronaldcutler/Library/CloudStorage//My Drive/Sidoli_Lab/Projects/Chromatin Decondensation/Data/RNA/Counts/Gencode/N3_2.genes.results" 
##                                                                                                                                                                       NaBut3_3 
##  "/Users/ronaldcutler/Library/CloudStorage//My Drive/Sidoli_Lab/Projects/Chromatin Decondensation/Data/RNA/Counts/Gencode/N3_3.genes.results" 
##                                                                                                                                                                      NaBut10_1 
## "/Users/ronaldcutler/Library/CloudStorage//My Drive/Sidoli_Lab/Projects/Chromatin Decondensation/Data/RNA/Counts/Gencode/N10_1.genes.results" 
##                                                                                                                                                                      NaBut10_2 
## "/Users/ronaldcutler/Library/CloudStorage//My Drive/Sidoli_Lab/Projects/Chromatin Decondensation/Data/RNA/Counts/Gencode/N10_2.genes.results" 
##                                                                                                                                                                      NaBut10_3 
## "/Users/ronaldcutler/Library/CloudStorage//My Drive/Sidoli_Lab/Projects/Chromatin Decondensation/Data/RNA/Counts/Gencode/N10_3.genes.results"
```

```
txi <- tximport(files, type="rsem", txIn = FALSE, txOut = FALSE, countsFromAbundance = "lengthScaledTPM")
```

```
## reading in files with read_tsv
```

```
## Warning in computeRsemGeneLevel(files, importer, geneIdCol, abundanceCol, :
## countsFromAbundance other than 'no' requires transcript-level estimates
```

```
## 1
```

```
## 2 3 4 5 6 7 8 9 10 11 12 13 14 15 16 17 18 19 20 21 22 23 24
```

```
se <- SummarizedExperiment(assays=SimpleList(tpm=txi$abundance,
                                             counts=txi$counts),
                                             colData=sampleTable)
rownames(se) <- lapply(rownames(se), sub, pattern = "\\.\\d+$", replacement = "") # remove version from gene name
```

Initial count distribution

```
plotDensity(log2(assay(se) + 1),
            lty=1,
            col=colors[as.factor(colData(se)$condition)],
            lwd=1,
            xlab = "log2(Counts + 1)", 
            main = "Raw Counts")
legend("topright",
       legend=levels(as.factor(colData(se)$condition)),
       lwd=1, col = colors)
```

```
boxplot(log2(assay(se) + 1),
        col=colors[as.factor(colData(se)$condition)],
        cex.axis = 0.5,
        las = 1,
        horizontal = TRUE,
        xlab = "log2(Counts + 1)",
        ylab = "Samples",
        main = "Raw Counts")
legend("topright",
       legend=levels(as.factor(colData(se)$condition)),
       lwd=1, col = colors)
```

Map ensemble names to gene names

```
ensembl <- useEnsembl(biomart = "genes", dataset = "hsapiens_gene_ensembl")

names <- getBM(attributes = c('ensembl_gene_id', 'external_gene_name', 'entrezgene_id', 'gene_biotype'),
                filters = 'ensembl_gene_id',
                values = rownames(se), 
                mart = ensembl)
```

Filter

```
assay(se, "zfpkm") <- zFPKM(se, assayName = "tpm")
assay(se, "zfpkm")[assay(se, "zfpkm") == -Inf] <- NA
zFPKMPlot(se, assayName = "tpm", FacetTitles = TRUE)
```

```
# To determine which genes are active, we compute the median expression within each group.
# detach(package:plyr)
activeGenes <- assay(se, "zfpkm") %>%
  mutate(gene=rownames(assay(se, "zfpkm"))) %>%
  gather(sample, zfpkm, -gene) %>%
  left_join(select(as.data.frame(colData(se)), sample, conditiontime), by="sample") %>%
  dplyr::group_by(gene, conditiontime) %>%
  summarize(median_zfpkm=median(zfpkm)) %>%
  ungroup() %>%
  mutate(active=(median_zfpkm > -3)) %>%
  filter(active) %>%
  select(gene) %>%
  distinct()
```

```
## `summarise()` has grouped output by 'gene'. You can override using the
## `.groups` argument.
```

```
se.filter <- SummarizedExperiment(
  assays=SimpleList(counts=as.matrix(round(assay(se, "counts")[activeGenes$gene, ]))),
  colData=colData(se))
```

Filtered count distribution

```
plotDensity(log2(assay(se.filter) + 1),
            lty=1,
            col=colors[as.factor(colData(se)$condition)],
            lwd=1,
            xlab = "log2(Counts + 1)", 
            main = "Filtered Counts")
legend("topright",
       legend=levels(as.factor(colData(se)$condition)),
       lwd=1, col = colors)
```

```
boxplot(log2(assay(se.filter) + 1),
        col=colors[as.factor(colData(se)$condition)],
        cex.axis = 0.5,
        las = 1,
        horizontal = TRUE,
        xlab = "log2(Counts + 1)",
        ylab = "Samples",
        main = "Filtered Counts")
legend("topright",
       legend=levels(as.factor(colData(se)$condition)),
       lwd=1, col = colors)
```

### Batch correction

```
# batch variable based off distance heatmap
mod <- model.matrix(~condition + time + condition:time, colData(se.filter))

se.filter.adjust <- ComBat_seq(assay(se.filter), batch = sampleTable$batch, covar_mod = mod)
```

```
## Found 2 batches
## Using null model in ComBat-seq.
## Adjusting for 7 covariate(s) or covariate level(s)
## Estimating dispersions
## Fitting the GLM model
## Shrinkage off - using GLM estimates for parameters
## Adjusting the data
```

Normalization

```
dds.adjust <- DESeqDataSetFromMatrix(se.filter.adjust, design = ~condition + time + condition:time, colData = sampleTable)
```

```
## converting counts to integer mode
```

```
## Warning in DESeqDataSet(se, design = design, ignoreRank): some variables in
## design formula are characters, converting to factors
```

```
dds.adjust <- DESeq(dds.adjust)
```

```
## estimating size factors
```

```
## estimating dispersions
```

```
## gene-wise dispersion estimates
```

```
## mean-dispersion relationship
```

```
## final dispersion estimates
```

```
## fitting model and testing
```

```
# normalized
dds.adjust.norm <- as.data.frame(counts(dds.adjust, normalized = TRUE))
dds.adjust.norm$symbol <- names[match(rownames(dds.adjust.norm), names$ensembl_gene_id), "external_gene_name"]
dds.adjust.norm$entrezgene_id <- names[match(rownames(dds.adjust.norm), names$ensembl_gene_id), "entrezgene_id"]
dds.adjust.norm$gene_biotype <- names[match(rownames(dds.adjust.norm), names$ensembl_gene_id), "gene_biotype"]

# VST
dds.adjust_vst <- vst(dds.adjust, blind = FALSE)
adjusted_vst <- as.data.frame(assay(dds.adjust_vst))
adjusted_vst$symbol <- names[match(rownames(adjusted_vst), names$ensembl_gene_id), "external_gene_name"]
adjusted_vst$entrezgene_id <- names[match(rownames(adjusted_vst), names$ensembl_gene_id), "entrezgene_id"]
adjusted_vst$gene_biotype <- names[match(rownames(adjusted_vst), names$ensembl_gene_id), "gene_biotype"]
```

Visualization

```
sampleDists <- dist(t(assay(dds.adjust_vst)))
sampleDistMatrix <- as.matrix(sampleDists)
rownames(sampleDistMatrix) <- paste(dds.adjust_vst$sample, dds.adjust_vst$conditiontime, sep="-")
colnames(sampleDistMatrix) <- paste(dds.adjust_vst$sample, dds.adjust_vst$conditiontime, sep="-")
colors <- colorRampPalette( rev(brewer.pal(9, "Blues")) )(255)
pheatmap(sampleDistMatrix,
         clustering_distance_rows=sampleDists,
         clustering_distance_cols=sampleDists,
         col=colors)
```

```
# set up matrix
dds_vst.mat <- as.matrix(assay(dds.adjust_vst))
select <- order(rowVars(dds_vst.mat), decreasing = TRUE)[1:(round(nrow(dds_vst.mat)*0.1))]
dds_vst.mat.var <- dds_vst.mat[select,]
ha <- HeatmapAnnotation(Condition = sampleTable$condition, Time = sampleTable$time)
scaled.mat <- t(scale(t(dds_vst.mat.var),center=TRUE,scale=TRUE))
Heatmap(scaled.mat, 
        name = "Z-score", #title of legend
        row_names_gp = gpar(fontsize = 7), # Text size for row names
        top_annotation = ha,
        border = TRUE,
        cluster_rows = TRUE,
        cluster_columns = TRUE,
        show_row_names = FALSE,
        column_names_rot = 45,
        show_parent_dend_line = FALSE,
        row_dend_width = unit(50, "mm"))
```

```
## `use_raster` is automatically set to TRUE for a matrix with more than
## 2000 rows. You can control `use_raster` argument by explicitly setting
## TRUE/FALSE to it.
## 
## Set `ht_opt$message = FALSE` to turn off this message.
```

```
umap_results <- umap(t(dds_vst.mat))
umap_plot_df <- data.frame(umap_results$layout) %>%
  tibble::rownames_to_column("sample") %>%
  dplyr::inner_join(sampleTable, by = "sample")
ggplot(umap_plot_df,aes(x = X1,y = X2)) +
  geom_point(aes(color = condition, shape = time), size = 3) +
  scale_color_viridis(discrete=TRUE) +
  theme_classic() +
  theme(axis.text.x=element_blank(),
      axis.text.y=element_blank(),
      axis.ticks=element_blank()) +
  xlab("UMAP 1") +
  ylab("UMAP 2")
```

MaSigPro regression models

```
edesign <- read.csv("/Users/ronaldcutler/Library/CloudStorage//My\ Drive/Sidoli_Lab/Projects/Chromatin\ Decondensation/Analysis/Gencode/masigpro.design.csv", header = TRUE, row.names = 1)

design <- make.design.matrix(edesign, degree = 3)

# find significant genes
fit <- p.vector(dds_vst.mat, design, Q = 0.05, MT.adjust = "BH", min.obs = 20, counts = TRUE)
```

```
## [1] "fitting  gene 100 out of 23655"
## [1] "fitting  gene 200 out of 23655"
## [1] "fitting  gene 300 out of 23655"
## [1] "fitting  gene 400 out of 23655"
## [1] "fitting  gene 500 out of 23655"
## [1] "fitting  gene 600 out of 23655"
## [1] "fitting  gene 700 out of 23655"
## [1] "fitting  gene 800 out of 23655"
## [1] "fitting  gene 900 out of 23655"
## [1] "fitting  gene 1000 out of 23655"
## [1] "fitting  gene 1100 out of 23655"
## [1] "fitting  gene 1200 out of 23655"
## [1] "fitting  gene 1300 out of 23655"
## [1] "fitting  gene 1400 out of 23655"
## [1] "fitting  gene 1500 out of 23655"
## [1] "fitting  gene 1600 out of 23655"
## [1] "fitting  gene 1700 out of 23655"
## [1] "fitting  gene 1800 out of 23655"
## [1] "fitting  gene 1900 out of 23655"
## [1] "fitting  gene 2000 out of 23655"
## [1] "fitting  gene 2100 out of 23655"
## [1] "fitting  gene 2200 out of 23655"
## [1] "fitting  gene 2300 out of 23655"
## [1] "fitting  gene 2400 out of 23655"
## [1] "fitting  gene 2500 out of 23655"
## [1] "fitting  gene 2600 out of 23655"
## [1] "fitting  gene 2700 out of 23655"
## [1] "fitting  gene 2800 out of 23655"
## [1] "fitting  gene 2900 out of 23655"
## [1] "fitting  gene 3000 out of 23655"
## [1] "fitting  gene 3100 out of 23655"
## [1] "fitting  gene 3200 out of 23655"
## [1] "fitting  gene 3300 out of 23655"
## [1] "fitting  gene 3400 out of 23655"
## [1] "fitting  gene 3500 out of 23655"
## [1] "fitting  gene 3600 out of 23655"
## [1] "fitting  gene 3700 out of 23655"
## [1] "fitting  gene 3800 out of 23655"
## [1] "fitting  gene 3900 out of 23655"
## [1] "fitting  gene 4000 out of 23655"
## [1] "fitting  gene 4100 out of 23655"
## [1] "fitting  gene 4200 out of 23655"
## [1] "fitting  gene 4300 out of 23655"
## [1] "fitting  gene 4400 out of 23655"
## [1] "fitting  gene 4500 out of 23655"
## [1] "fitting  gene 4600 out of 23655"
## [1] "fitting  gene 4700 out of 23655"
## [1] "fitting  gene 4800 out of 23655"
## [1] "fitting  gene 4900 out of 23655"
## [1] "fitting  gene 5000 out of 23655"
## [1] "fitting  gene 5100 out of 23655"
## [1] "fitting  gene 5200 out of 23655"
## [1] "fitting  gene 5300 out of 23655"
## [1] "fitting  gene 5400 out of 23655"
## [1] "fitting  gene 5500 out of 23655"
## [1] "fitting  gene 5600 out of 23655"
## [1] "fitting  gene 5700 out of 23655"
## [1] "fitting  gene 5800 out of 23655"
## [1] "fitting  gene 5900 out of 23655"
## [1] "fitting  gene 6000 out of 23655"
## [1] "fitting  gene 6100 out of 23655"
## [1] "fitting  gene 6200 out of 23655"
## [1] "fitting  gene 6300 out of 23655"
## [1] "fitting  gene 6400 out of 23655"
## [1] "fitting  gene 6500 out of 23655"
## [1] "fitting  gene 6600 out of 23655"
## [1] "fitting  gene 6700 out of 23655"
## [1] "fitting  gene 6800 out of 23655"
## [1] "fitting  gene 6900 out of 23655"
## [1] "fitting  gene 7000 out of 23655"
## [1] "fitting  gene 7100 out of 23655"
## [1] "fitting  gene 7200 out of 23655"
## [1] "fitting  gene 7300 out of 23655"
## [1] "fitting  gene 7400 out of 23655"
## [1] "fitting  gene 7500 out of 23655"
## [1] "fitting  gene 7600 out of 23655"
## [1] "fitting  gene 7700 out of 23655"
## [1] "fitting  gene 7800 out of 23655"
## [1] "fitting  gene 7900 out of 23655"
## [1] "fitting  gene 8000 out of 23655"
## [1] "fitting  gene 8100 out of 23655"
## [1] "fitting  gene 8200 out of 23655"
## [1] "fitting  gene 8300 out of 23655"
## [1] "fitting  gene 8400 out of 23655"
## [1] "fitting  gene 8500 out of 23655"
## [1] "fitting  gene 8600 out of 23655"
## [1] "fitting  gene 8700 out of 23655"
## [1] "fitting  gene 8800 out of 23655"
## [1] "fitting  gene 8900 out of 23655"
## [1] "fitting  gene 9000 out of 23655"
## [1] "fitting  gene 9100 out of 23655"
## [1] "fitting  gene 9200 out of 23655"
## [1] "fitting  gene 9300 out of 23655"
## [1] "fitting  gene 9400 out of 23655"
## [1] "fitting  gene 9500 out of 23655"
## [1] "fitting  gene 9600 out of 23655"
## [1] "fitting  gene 9700 out of 23655"
## [1] "fitting  gene 9800 out of 23655"
## [1] "fitting  gene 9900 out of 23655"
## [1] "fitting  gene 10000 out of 23655"
## [1] "fitting  gene 10100 out of 23655"
## [1] "fitting  gene 10200 out of 23655"
## [1] "fitting  gene 10300 out of 23655"
## [1] "fitting  gene 10400 out of 23655"
## [1] "fitting  gene 10500 out of 23655"
## [1] "fitting  gene 10600 out of 23655"
## [1] "fitting  gene 10700 out of 23655"
## [1] "fitting  gene 10800 out of 23655"
## [1] "fitting  gene 10900 out of 23655"
## [1] "fitting  gene 11000 out of 23655"
## [1] "fitting  gene 11100 out of 23655"
## [1] "fitting  gene 11200 out of 23655"
## [1] "fitting  gene 11300 out of 23655"
## [1] "fitting  gene 11400 out of 23655"
## [1] "fitting  gene 11500 out of 23655"
## [1] "fitting  gene 11600 out of 23655"
## [1] "fitting  gene 11700 out of 23655"
## [1] "fitting  gene 11800 out of 23655"
## [1] "fitting  gene 11900 out of 23655"
## [1] "fitting  gene 12000 out of 23655"
## [1] "fitting  gene 12100 out of 23655"
## [1] "fitting  gene 12200 out of 23655"
## [1] "fitting  gene 12300 out of 23655"
## [1] "fitting  gene 12400 out of 23655"
## [1] "fitting  gene 12500 out of 23655"
## [1] "fitting  gene 12600 out of 23655"
## [1] "fitting  gene 12700 out of 23655"
## [1] "fitting  gene 12800 out of 23655"
## [1] "fitting  gene 12900 out of 23655"
## [1] "fitting  gene 13000 out of 23655"
## [1] "fitting  gene 13100 out of 23655"
## [1] "fitting  gene 13200 out of 23655"
## [1] "fitting  gene 13300 out of 23655"
## [1] "fitting  gene 13400 out of 23655"
## [1] "fitting  gene 13500 out of 23655"
## [1] "fitting  gene 13600 out of 23655"
## [1] "fitting  gene 13700 out of 23655"
## [1] "fitting  gene 13800 out of 23655"
## [1] "fitting  gene 13900 out of 23655"
## [1] "fitting  gene 14000 out of 23655"
## [1] "fitting  gene 14100 out of 23655"
## [1] "fitting  gene 14200 out of 23655"
## [1] "fitting  gene 14300 out of 23655"
## [1] "fitting  gene 14400 out of 23655"
## [1] "fitting  gene 14500 out of 23655"
## [1] "fitting  gene 14600 out of 23655"
## [1] "fitting  gene 14700 out of 23655"
## [1] "fitting  gene 14800 out of 23655"
## [1] "fitting  gene 14900 out of 23655"
## [1] "fitting  gene 15000 out of 23655"
## [1] "fitting  gene 15100 out of 23655"
## [1] "fitting  gene 15200 out of 23655"
## [1] "fitting  gene 15300 out of 23655"
## [1] "fitting  gene 15400 out of 23655"
## [1] "fitting  gene 15500 out of 23655"
## [1] "fitting  gene 15600 out of 23655"
## [1] "fitting  gene 15700 out of 23655"
## [1] "fitting  gene 15800 out of 23655"
## [1] "fitting  gene 15900 out of 23655"
## [1] "fitting  gene 16000 out of 23655"
## [1] "fitting  gene 16100 out of 23655"
## [1] "fitting  gene 16200 out of 23655"
## [1] "fitting  gene 16300 out of 23655"
## [1] "fitting  gene 16400 out of 23655"
## [1] "fitting  gene 16500 out of 23655"
## [1] "fitting  gene 16600 out of 23655"
## [1] "fitting  gene 16700 out of 23655"
## [1] "fitting  gene 16800 out of 23655"
## [1] "fitting  gene 16900 out of 23655"
## [1] "fitting  gene 17000 out of 23655"
## [1] "fitting  gene 17100 out of 23655"
## [1] "fitting  gene 17200 out of 23655"
## [1] "fitting  gene 17300 out of 23655"
## [1] "fitting  gene 17400 out of 23655"
## [1] "fitting  gene 17500 out of 23655"
## [1] "fitting  gene 17600 out of 23655"
## [1] "fitting  gene 17700 out of 23655"
## [1] "fitting  gene 17800 out of 23655"
## [1] "fitting  gene 17900 out of 23655"
## [1] "fitting  gene 18000 out of 23655"
## [1] "fitting  gene 18100 out of 23655"
## [1] "fitting  gene 18200 out of 23655"
## [1] "fitting  gene 18300 out of 23655"
## [1] "fitting  gene 18400 out of 23655"
## [1] "fitting  gene 18500 out of 23655"
## [1] "fitting  gene 18600 out of 23655"
## [1] "fitting  gene 18700 out of 23655"
## [1] "fitting  gene 18800 out of 23655"
## [1] "fitting  gene 18900 out of 23655"
## [1] "fitting  gene 19000 out of 23655"
## [1] "fitting  gene 19100 out of 23655"
## [1] "fitting  gene 19200 out of 23655"
## [1] "fitting  gene 19300 out of 23655"
## [1] "fitting  gene 19400 out of 23655"
## [1] "fitting  gene 19500 out of 23655"
## [1] "fitting  gene 19600 out of 23655"
## [1] "fitting  gene 19700 out of 23655"
## [1] "fitting  gene 19800 out of 23655"
## [1] "fitting  gene 19900 out of 23655"
## [1] "fitting  gene 20000 out of 23655"
## [1] "fitting  gene 20100 out of 23655"
## [1] "fitting  gene 20200 out of 23655"
## [1] "fitting  gene 20300 out of 23655"
## [1] "fitting  gene 20400 out of 23655"
## [1] "fitting  gene 20500 out of 23655"
## [1] "fitting  gene 20600 out of 23655"
## [1] "fitting  gene 20700 out of 23655"
## [1] "fitting  gene 20800 out of 23655"
## [1] "fitting  gene 20900 out of 23655"
## [1] "fitting  gene 21000 out of 23655"
## [1] "fitting  gene 21100 out of 23655"
## [1] "fitting  gene 21200 out of 23655"
## [1] "fitting  gene 21300 out of 23655"
## [1] "fitting  gene 21400 out of 23655"
## [1] "fitting  gene 21500 out of 23655"
## [1] "fitting  gene 21600 out of 23655"
## [1] "fitting  gene 21700 out of 23655"
## [1] "fitting  gene 21800 out of 23655"
## [1] "fitting  gene 21900 out of 23655"
## [1] "fitting  gene 22000 out of 23655"
## [1] "fitting  gene 22100 out of 23655"
## [1] "fitting  gene 22200 out of 23655"
## [1] "fitting  gene 22300 out of 23655"
## [1] "fitting  gene 22400 out of 23655"
## [1] "fitting  gene 22500 out of 23655"
## [1] "fitting  gene 22600 out of 23655"
## [1] "fitting  gene 22700 out of 23655"
## [1] "fitting  gene 22800 out of 23655"
## [1] "fitting  gene 22900 out of 23655"
## [1] "fitting  gene 23000 out of 23655"
## [1] "fitting  gene 23100 out of 23655"
## [1] "fitting  gene 23200 out of 23655"
## [1] "fitting  gene 23300 out of 23655"
## [1] "fitting  gene 23400 out of 23655"
## [1] "fitting  gene 23500 out of 23655"
## [1] "fitting  gene 23600 out of 23655"
```

```
# find significant differences
tstep <- T.fit(fit, step.method = "backward", alfa = 0.05)
```

```
## [1] "fitting  gene 100 out of 13437"
## [1] "fitting  gene 200 out of 13437"
## [1] "fitting  gene 300 out of 13437"
## [1] "fitting  gene 400 out of 13437"
## [1] "fitting  gene 500 out of 13437"
## [1] "fitting  gene 600 out of 13437"
## [1] "fitting  gene 700 out of 13437"
## [1] "fitting  gene 800 out of 13437"
## [1] "fitting  gene 900 out of 13437"
## [1] "fitting  gene 1000 out of 13437"
## [1] "fitting  gene 1100 out of 13437"
## [1] "fitting  gene 1200 out of 13437"
## [1] "fitting  gene 1300 out of 13437"
## [1] "fitting  gene 1400 out of 13437"
## [1] "fitting  gene 1500 out of 13437"
## [1] "fitting  gene 1600 out of 13437"
## [1] "fitting  gene 1700 out of 13437"
## [1] "fitting  gene 1800 out of 13437"
## [1] "fitting  gene 1900 out of 13437"
## [1] "fitting  gene 2000 out of 13437"
## [1] "fitting  gene 2100 out of 13437"
## [1] "fitting  gene 2200 out of 13437"
## [1] "fitting  gene 2300 out of 13437"
## [1] "fitting  gene 2400 out of 13437"
## [1] "fitting  gene 2500 out of 13437"
## [1] "fitting  gene 2600 out of 13437"
## [1] "fitting  gene 2700 out of 13437"
## [1] "fitting  gene 2800 out of 13437"
## [1] "fitting  gene 2900 out of 13437"
## [1] "fitting  gene 3000 out of 13437"
## [1] "fitting  gene 3100 out of 13437"
## [1] "fitting  gene 3200 out of 13437"
## [1] "fitting  gene 3300 out of 13437"
## [1] "fitting  gene 3400 out of 13437"
## [1] "fitting  gene 3500 out of 13437"
## [1] "fitting  gene 3600 out of 13437"
## [1] "fitting  gene 3700 out of 13437"
## [1] "fitting  gene 3800 out of 13437"
## [1] "fitting  gene 3900 out of 13437"
## [1] "fitting  gene 4000 out of 13437"
## [1] "fitting  gene 4100 out of 13437"
## [1] "fitting  gene 4200 out of 13437"
## [1] "fitting  gene 4300 out of 13437"
## [1] "fitting  gene 4400 out of 13437"
## [1] "fitting  gene 4500 out of 13437"
## [1] "fitting  gene 4600 out of 13437"
## [1] "fitting  gene 4700 out of 13437"
## [1] "fitting  gene 4800 out of 13437"
## [1] "fitting  gene 4900 out of 13437"
## [1] "fitting  gene 5000 out of 13437"
## [1] "fitting  gene 5100 out of 13437"
## [1] "fitting  gene 5200 out of 13437"
## [1] "fitting  gene 5300 out of 13437"
## [1] "fitting  gene 5400 out of 13437"
## [1] "fitting  gene 5500 out of 13437"
## [1] "fitting  gene 5600 out of 13437"
## [1] "fitting  gene 5700 out of 13437"
## [1] "fitting  gene 5800 out of 13437"
## [1] "fitting  gene 5900 out of 13437"
## [1] "fitting  gene 6000 out of 13437"
## [1] "fitting  gene 6100 out of 13437"
## [1] "fitting  gene 6200 out of 13437"
## [1] "fitting  gene 6300 out of 13437"
## [1] "fitting  gene 6400 out of 13437"
## [1] "fitting  gene 6500 out of 13437"
## [1] "fitting  gene 6600 out of 13437"
## [1] "fitting  gene 6700 out of 13437"
## [1] "fitting  gene 6800 out of 13437"
## [1] "fitting  gene 6900 out of 13437"
## [1] "fitting  gene 7000 out of 13437"
## [1] "fitting  gene 7100 out of 13437"
## [1] "fitting  gene 7200 out of 13437"
## [1] "fitting  gene 7300 out of 13437"
## [1] "fitting  gene 7400 out of 13437"
## [1] "fitting  gene 7500 out of 13437"
## [1] "fitting  gene 7600 out of 13437"
## [1] "fitting  gene 7700 out of 13437"
## [1] "fitting  gene 7800 out of 13437"
## [1] "fitting  gene 7900 out of 13437"
## [1] "fitting  gene 8000 out of 13437"
## [1] "fitting  gene 8100 out of 13437"
## [1] "fitting  gene 8200 out of 13437"
## [1] "fitting  gene 8300 out of 13437"
## [1] "fitting  gene 8400 out of 13437"
## [1] "fitting  gene 8500 out of 13437"
## [1] "fitting  gene 8600 out of 13437"
## [1] "fitting  gene 8700 out of 13437"
## [1] "fitting  gene 8800 out of 13437"
## [1] "fitting  gene 8900 out of 13437"
## [1] "fitting  gene 9000 out of 13437"
## [1] "fitting  gene 9100 out of 13437"
## [1] "fitting  gene 9200 out of 13437"
## [1] "fitting  gene 9300 out of 13437"
## [1] "fitting  gene 9400 out of 13437"
## [1] "fitting  gene 9500 out of 13437"
## [1] "fitting  gene 9600 out of 13437"
## [1] "fitting  gene 9700 out of 13437"
## [1] "fitting  gene 9800 out of 13437"
## [1] "fitting  gene 9900 out of 13437"
## [1] "fitting  gene 10000 out of 13437"
## [1] "fitting  gene 10100 out of 13437"
## [1] "fitting  gene 10200 out of 13437"
## [1] "fitting  gene 10300 out of 13437"
## [1] "fitting  gene 10400 out of 13437"
## [1] "fitting  gene 10500 out of 13437"
## [1] "fitting  gene 10600 out of 13437"
## [1] "fitting  gene 10700 out of 13437"
## [1] "fitting  gene 10800 out of 13437"
## [1] "fitting  gene 10900 out of 13437"
## [1] "fitting  gene 11000 out of 13437"
## [1] "fitting  gene 11100 out of 13437"
## [1] "fitting  gene 11200 out of 13437"
## [1] "fitting  gene 11300 out of 13437"
## [1] "fitting  gene 11400 out of 13437"
## [1] "fitting  gene 11500 out of 13437"
## [1] "fitting  gene 11600 out of 13437"
## [1] "fitting  gene 11700 out of 13437"
## [1] "fitting  gene 11800 out of 13437"
## [1] "fitting  gene 11900 out of 13437"
## [1] "fitting  gene 12000 out of 13437"
## [1] "fitting  gene 12100 out of 13437"
## [1] "fitting  gene 12200 out of 13437"
## [1] "fitting  gene 12300 out of 13437"
## [1] "fitting  gene 12400 out of 13437"
## [1] "fitting  gene 12500 out of 13437"
## [1] "fitting  gene 12600 out of 13437"
## [1] "fitting  gene 12700 out of 13437"
## [1] "fitting  gene 12800 out of 13437"
## [1] "fitting  gene 12900 out of 13437"
## [1] "fitting  gene 13000 out of 13437"
## [1] "fitting  gene 13100 out of 13437"
## [1] "fitting  gene 13200 out of 13437"
## [1] "fitting  gene 13300 out of 13437"
## [1] "fitting  gene 13400 out of 13437"
## [1] "Influence: 2655 genes with influential data at slot influ.info. Model validation for these genes is recommended"
```

```
# obtain list of significant genes
sigs <- get.siggenes(tstep, rsq = 0.7, vars = "groups")
```

Control only - Genes that change over the course of the experiment - 1719 genes

```
# number of significant
sigs$sig.genes$Control$g
```

```
## [1] 1762
```

```
# plot these genes 
library(mclust)
```

```
## Package 'mclust' version 5.4.9
## Type 'citation("mclust")' for citing this R package in publications.
```

```
## 
## Attaching package: 'mclust'
```

```
## The following object is masked from 'package:purrr':
## 
##     map
```

```
## The following object is masked from 'package:mgcv':
## 
##     mvn
```

```
see.genes(sigs$sig.genes$Control, show.fit = T, dis =design$dis, cluster.method="Mclust", cluster.data = 1, k.mclust=TRUE)
```

```
## $cut
##          ENSG00000001630          ENSG00000004478          ENSG00000005187 
##                        1                        1                        7 
##          ENSG00000005189          ENSG00000005486          ENSG00000005884 
##                        3                        8                        2 
##          ENSG00000005889          ENSG00000005961          ENSG00000006047 
##                        1                        7                        4 
##          ENSG00000007080          ENSG00000007516          ENSG00000007944 
##                        2                        2                        7 
##          ENSG00000008735          ENSG00000009950          ENSG00000010278 
##                        8                        8                        2 
##          ENSG00000010295          ENSG00000010327          ENSG00000011347 
##                        8                        2                        2 
##          ENSG00000011638          ENSG00000012124          ENSG00000012171 
##                        4                        2                        1 
##          ENSG00000013588          ENSG00000013725          ENSG00000016391 
##                        7                        6                        8 
##          ENSG00000017483          ENSG00000018699          ENSG00000019102 
##                        1                        5                        5 
##          ENSG00000019186          ENSG00000021355          ENSG00000023445 
##                        7                        8                        7 
##          ENSG00000023734          ENSG00000023909          ENSG00000026103 
##                        8                        7                        5 
##          ENSG00000030110          ENSG00000030582          ENSG00000035862 
##                        8                        8                        8 
##          ENSG00000037042          ENSG00000037280          ENSG00000039068 
##                        2                        6                        8 
##          ENSG00000039987          ENSG00000040608          ENSG00000042781 
##                        6                        4                        7 
##          ENSG00000044446          ENSG00000044524          ENSG00000049249 
##                        5                        6                        4 
##          ENSG00000049449          ENSG00000049540          ENSG00000050405 
##                        8                        4                        2 
##          ENSG00000050555          ENSG00000051108          ENSG00000052344 
##                        6                        8                        2 
##          ENSG00000052795          ENSG00000052802          ENSG00000053254 
##                        8                        1                        8 
##          ENSG00000054598          ENSG00000054654          ENSG00000054793 
##                        2                        7                        7 
##          ENSG00000055163          ENSG00000055957          ENSG00000056558 
##                        8                        5                        4 
##          ENSG00000056998          ENSG00000057593          ENSG00000057704 
##                        1                        5                        6 
##          ENSG00000059122          ENSG00000059728          ENSG00000059804 
##                        8                        8                        8 
##          ENSG00000059915          ENSG00000060558          ENSG00000060642 
##                        2                        6                        8 
##          ENSG00000061337          ENSG00000062282          ENSG00000062524 
##                        6                        8                        2 
##          ENSG00000065361          ENSG00000065675          ENSG00000065717 
##                        1                        6                        2 
##          ENSG00000065911          ENSG00000066056          ENSG00000066583 
##                        1                        7                        8 
##          ENSG00000066813          ENSG00000067064          ENSG00000067842 
##                        5                        8                        6 
##          ENSG00000068400          ENSG00000068971          ENSG00000069011 
##                        8                        8                        1 
##          ENSG00000069812          ENSG00000069849          ENSG00000070495 
##                        4                        8                        8 
##          ENSG00000070614          ENSG00000070729          ENSG00000070915 
##                        8                        4                        5 
##          ENSG00000071859          ENSG00000071909          ENSG00000072163 
##                        8                        3                        2 
##          ENSG00000072195          ENSG00000072401          ENSG00000072422 
##                        4                        2                        5 
##          ENSG00000072954          ENSG00000073150          ENSG00000073331 
##                        2                        5                        2 
##          ENSG00000073670          ENSG00000074181          ENSG00000074410 
##                        2                        2                        4 
##          ENSG00000075223          ENSG00000075340          ENSG00000075618 
##                        4                        6                        8 
##          ENSG00000076067          ENSG00000076258          ENSG00000076604 
##                        2                        6                        1 
##          ENSG00000076706          ENSG00000076770          ENSG00000077782 
##                        2                        8                        8 
##          ENSG00000078081          ENSG00000078124          ENSG00000078898 
##                        4                        7                        8 
##          ENSG00000079459          ENSG00000079999          ENSG00000080561 
##                        8                        1                        6 
##          ENSG00000080573          ENSG00000081051          ENSG00000081479 
##                        4                        8                        7 
##          ENSG00000081923          ENSG00000082074          ENSG00000082512 
##                        2                        6                        4 
##          ENSG00000082684          ENSG00000083828          ENSG00000084110 
##                        3                        3                        8 
##          ENSG00000084710          ENSG00000084734          ENSG00000085552 
##                        2                        5                        6 
##          ENSG00000085662          ENSG00000086159          ENSG00000086696 
##                        8                        2                        5 
##          ENSG00000087074          ENSG00000087076          ENSG00000088387 
##                        8                        6                        4 
##          ENSG00000088826          ENSG00000089060          ENSG00000089116 
##                        8                        2                        6 
##          ENSG00000089127          ENSG00000089199          ENSG00000089356 
##                        2                        4                        5 
##          ENSG00000090020          ENSG00000090238          ENSG00000090382 
##                        8                        2                        1 
##          ENSG00000090447          ENSG00000090512          ENSG00000090520 
##                        5                        5                        8 
##          ENSG00000090612          ENSG00000091106          ENSG00000091262 
##                        2                        3                        1 
##          ENSG00000091513          ENSG00000091536          ENSG00000091844 
##                        8                        4                        7 
##          ENSG00000092607          ENSG00000092621          ENSG00000092758 
##                        5                        1                        8 
##          ENSG00000092820          ENSG00000095383          ENSG00000095485 
##                        8                        2                        1 
##          ENSG00000095539          ENSG00000095739          ENSG00000095752 
##                        8                        8                        7 
##          ENSG00000096060          ENSG00000096088          ENSG00000099139 
##                        8                        8                        4 
##          ENSG00000099250          ENSG00000099251          ENSG00000099282 
##                        5                        5                        2 
##          ENSG00000099284          ENSG00000099797          ENSG00000099812 
##                        5                        8                        8 
##          ENSG00000099840          ENSG00000099860          ENSG00000099953 
##                        8                        8                        2 
##          ENSG00000099985          ENSG00000099994          ENSG00000100239 
##                        6                        8                        1 
##          ENSG00000100290          ENSG00000100300          ENSG00000100307 
##                        7                        2                        2 
##          ENSG00000100344          ENSG00000100373          ENSG00000100385 
##                        1                        5                        6 
##          ENSG00000100399          ENSG00000100427          ENSG00000100554 
##                        3                        6                        2 
##          ENSG00000100600          ENSG00000100665          ENSG00000100867 
##                        8                        8                        8 
##          ENSG00000100889          ENSG00000100985          ENSG00000101000 
##                        1                        4                        4 
##          ENSG00000101162          ENSG00000101187          ENSG00000101188 
##                        7                        1                        3 
##          ENSG00000101197          ENSG00000101210          ENSG00000101265 
##                        8                        2                        4 
##          ENSG00000101276          ENSG00000101342          ENSG00000101353 
##                        4                        2                        3 
##          ENSG00000101384          ENSG00000101670          ENSG00000101846 
##                        1                        7                        5 
##          ENSG00000102003          ENSG00000102048          ENSG00000102109 
##                        2                        5                        2 
##          ENSG00000102265          ENSG00000102554          ENSG00000102760 
##                        7                        2                        4 
##          ENSG00000103034          ENSG00000103154          ENSG00000103160 
##                        7                        6                        8 
##          ENSG00000103187          ENSG00000103196          ENSG00000103313 
##                        8                        7                        6 
##          ENSG00000103316          ENSG00000103449          ENSG00000103710 
##                        2                        4                        6 
##          ENSG00000103742          ENSG00000103888          ENSG00000104081 
##                        4                        5                        8 
##          ENSG00000104408          ENSG00000104549          ENSG00000104626 
##                        1                        8                        2 
##          ENSG00000104635          ENSG00000104814          ENSG00000104889 
##                        1                        4                        1 
##          ENSG00000104892          ENSG00000104907          ENSG00000104969 
##                        2                        1                        8 
##          ENSG00000104979          ENSG00000105088          ENSG00000105135 
##                        7                        4                        1 
##          ENSG00000105227          ENSG00000105255          ENSG00000105327 
##                        2                        4                        8 
##          ENSG00000105339          ENSG00000105370          ENSG00000105398 
##                        2                        5                        8 
##          ENSG00000105401          ENSG00000105419          ENSG00000105538 
##                        8                        2                        2 
##          ENSG00000105642          ENSG00000105664          ENSG00000105668 
##                        4                        2                        6 
##          ENSG00000105696          ENSG00000105697          ENSG00000105707 
##                        2                        2                        8 
##          ENSG00000105737          ENSG00000105784          ENSG00000105808 
##                        6                        3                        7 
##          ENSG00000106327          ENSG00000106415          ENSG00000106538 
##                        8                        2                        2 
##          ENSG00000106624          ENSG00000106780          ENSG00000106992 
##                        3                        8                        2 
##          ENSG00000107186          ENSG00000107331          ENSG00000107731 
##                        6                        8                        7 
##          ENSG00000107859          ENSG00000107984          ENSG00000108106 
##                        4                        7                        8 
##          ENSG00000108244          ENSG00000108309          ENSG00000108379 
##                        7                        7                        5 
##          ENSG00000108518          ENSG00000108551          ENSG00000108561 
##                        8                        5                        8 
##          ENSG00000108576          ENSG00000108846          ENSG00000108932 
##                        1                        5                        5 
##          ENSG00000108984          ENSG00000109072          ENSG00000109084 
##                        4                        8                        1 
##          ENSG00000109089          ENSG00000109113          ENSG00000109181 
##                        2                        5                        5 
##          ENSG00000109193          ENSG00000109321          ENSG00000109501 
##                        7                        4                        7 
##          ENSG00000109667          ENSG00000109680          ENSG00000109758 
##                        1                        4                        1 
##          ENSG00000109771          ENSG00000109971          ENSG00000110046 
##                        3                        8                        8 
##          ENSG00000110169          ENSG00000110237          ENSG00000110243 
##                        7                        1                        5 
##          ENSG00000110697          ENSG00000110799          ENSG00000110911 
##                        8                        5                        1 
##          ENSG00000110921          ENSG00000110934          ENSG00000111186 
##                        8                        6                        6 
##          ENSG00000111271          ENSG00000111339          ENSG00000111348 
##                        8                        7                        6 
##          ENSG00000111452          ENSG00000111679          ENSG00000111788 
##                        5                        2                        5 
##          ENSG00000111860          ENSG00000111879          ENSG00000112079 
##                        4                        7                        2 
##          ENSG00000112139          ENSG00000112182          ENSG00000112183 
##                        6                        4                        5 
##          ENSG00000112406          ENSG00000112494          ENSG00000112561 
##                        4                        5                        2 
##          ENSG00000112576          ENSG00000112773          ENSG00000112964 
##                        8                        8                        4 
##          ENSG00000112972          ENSG00000112992          ENSG00000113119 
##                        8                        1                        2 
##          ENSG00000113141          ENSG00000113161          ENSG00000113212 
##                        8                        8                        6 
##          ENSG00000113273          ENSG00000113312          ENSG00000113319 
##                        7                        2                        5 
##          ENSG00000113396          ENSG00000113504          ENSG00000113790 
##                        6                        1                        1 
##          ENSG00000113924          ENSG00000113966          ENSG00000114541 
##                        1                        4                        2 
##          ENSG00000114739          ENSG00000114933          ENSG00000114999 
##                        1                        7                        8 
##          ENSG00000115112          ENSG00000115257          ENSG00000115267 
##                        6                        2                        2 
##          ENSG00000115271          ENSG00000115310          ENSG00000115468 
##                        2                        8                        2 
##          ENSG00000115758          ENSG00000116001          ENSG00000116014 
##                        8                        5                        8 
##          ENSG00000116133          ENSG00000116254          ENSG00000116329 
##                        8                        2                        4 
##          ENSG00000116455          ENSG00000116663          ENSG00000116667 
##                        5                        5                        6 
##          ENSG00000116690          ENSG00000116704          ENSG00000116717 
##                        3                        1                        8 
##          ENSG00000116833          ENSG00000116852          ENSG00000116871 
##                        8                        7                        8 
##          ENSG00000116885          ENSG00000116990          ENSG00000117280 
##                        6                        2                        2 
##          ENSG00000117318          ENSG00000117399          ENSG00000117601 
##                        8                        1                        8 
##          ENSG00000117643          ENSG00000117707          ENSG00000117791 
##                        4                        8                        2 
##          ENSG00000117834          ENSG00000117983          ENSG00000118137 
##                        8                        3                        8 
##          ENSG00000118271          ENSG00000118298          ENSG00000118520 
##                        7                        3                        3 
##          ENSG00000118523          ENSG00000118690          ENSG00000118785 
##                        8                        4                        7 
##          ENSG00000118898          ENSG00000118971          ENSG00000119138 
##                        2                        8                        2 
##          ENSG00000119227          ENSG00000119411          ENSG00000119535 
##                        5                        2                        5 
##          ENSG00000119673          ENSG00000119711          ENSG00000119723 
##                        1                        1                        7 
##          ENSG00000119729          ENSG00000119917          ENSG00000119927 
##                        7                        3                        1 
##          ENSG00000119938          ENSG00000120129          ENSG00000120322 
##                        2                        8                        4 
##          ENSG00000120327          ENSG00000120341          ENSG00000120694 
##                        3                        5                        8 
##          ENSG00000120738          ENSG00000120784          ENSG00000120875 
##                        8                        5                        4 
##          ENSG00000120885          ENSG00000120903          ENSG00000120913 
##                        8                        2                        1 
##          ENSG00000120937          ENSG00000121211          ENSG00000121310 
##                        3                        7                        5 
##          ENSG00000121410          ENSG00000121653          ENSG00000121680 
##                        5                        2                        2 
##          ENSG00000121716          ENSG00000121769          ENSG00000121858 
##                        1                        3                        5 
##          ENSG00000122121          ENSG00000122133          ENSG00000122194 
##                        5                        4                        3 
##          ENSG00000122490          ENSG00000122787          ENSG00000122861 
##                        8                        5                        4 
##          ENSG00000123095          ENSG00000123096          ENSG00000123146 
##                        6                        6                        2 
##          ENSG00000123358          ENSG00000123453          ENSG00000123600 
##                        8                        1                        1 
##          ENSG00000123636          ENSG00000123843          ENSG00000123933 
##                        5                        5                        8 
##          ENSG00000123999          ENSG00000124092          ENSG00000124104 
##                        5                        4                        2 
##          ENSG00000124216          ENSG00000124243          ENSG00000124253 
##                        2                        2                        8 
##          ENSG00000124557          ENSG00000124568          ENSG00000124713 
##                        4                        3                        2 
##          ENSG00000124731          ENSG00000124762          ENSG00000125144 
##                        4                        8                        8 
##          ENSG00000125148          ENSG00000125257          ENSG00000125378 
##                        8                        7                        8 
##          ENSG00000125508          ENSG00000125657          ENSG00000125703 
##                        2                        2                        2 
##          ENSG00000125731          ENSG00000125735          ENSG00000125740 
##                        7                        5                        2 
##          ENSG00000125743          ENSG00000125744          ENSG00000125775 
##                        8                        2                        5 
##          ENSG00000126091          ENSG00000126217          ENSG00000126231 
##                        5                        8                        3 
##          ENSG00000126259          ENSG00000126464          ENSG00000126561 
##                        4                        8                        5 
##          ENSG00000126583          ENSG00000126709          ENSG00000127241 
##                        6                        7                        8 
##          ENSG00000127324          ENSG00000127507          ENSG00000127561 
##                        3                        3                        2 
##          ENSG00000127578          ENSG00000127585          ENSG00000127743 
##                        5                        2                        5 
##          ENSG00000128011          ENSG00000128016          ENSG00000128039 
##                        4                        7                        2 
##          ENSG00000128268          ENSG00000128311          ENSG00000128342 
##                        3                        8                        5 
##          ENSG00000128394          ENSG00000128482          ENSG00000128487 
##                        2                        6                        2 
##          ENSG00000128564          ENSG00000128602          ENSG00000128656 
##                        7                        1                        4 
##          ENSG00000128872          ENSG00000128965          ENSG00000129038 
##                        4                        4                        6 
##          ENSG00000129214          ENSG00000129219          ENSG00000129244 
##                        2                        8                        7 
##          ENSG00000129465          ENSG00000129474          ENSG00000129538 
##                        6                        1                        6 
##          ENSG00000129596          ENSG00000129910          ENSG00000129946 
##                        2                        2                        5 
##          ENSG00000129991          ENSG00000130023          ENSG00000130066 
##                        4                        2                        8 
##          ENSG00000130158          ENSG00000130165          ENSG00000130173 
##                        1                        8                        8 
##          ENSG00000130270          ENSG00000130300          ENSG00000130349 
##                        2                        6                        2 
##          ENSG00000130427          ENSG00000130479          ENSG00000130487 
##                        5                        8                        5 
##          ENSG00000130600          ENSG00000130818          ENSG00000130822 
##                        2                        1                        8 
##          ENSG00000130844          ENSG00000130956          ENSG00000130957 
##                        5                        2                        4 
##          ENSG00000130988          ENSG00000131037          ENSG00000131069 
##                        5                        2                        1 
##          ENSG00000131094          ENSG00000131095          ENSG00000131187 
##                        4                        4                        5 
##          ENSG00000131482          ENSG00000131650          ENSG00000131711 
##                        5                        7                        8 
##          ENSG00000131730          ENSG00000131797          ENSG00000131876 
##                        5                        1                        8 
##          ENSG00000131899          ENSG00000131981          ENSG00000132002 
##                        8                        8                        8 
##          ENSG00000132026          ENSG00000132196          ENSG00000132437 
##                        2                        1                        1 
##          ENSG00000132669          ENSG00000132677          ENSG00000132688 
##                        5                        8                        2 
##          ENSG00000132698          ENSG00000132718          ENSG00000132740 
##                        2                        2                        5 
##          ENSG00000132768          ENSG00000132855          ENSG00000132881 
##                        1                        5                        2 
##          ENSG00000133169          ENSG00000133316          ENSG00000133640 
##                        4                        5                        6 
##          ENSG00000133665          ENSG00000133794          ENSG00000133943 
##                        4                        2                        8 
##          ENSG00000134013          ENSG00000134240          ENSG00000134297 
##                        2                        7                        2 
##          ENSG00000134470          ENSG00000134824          ENSG00000134853 
##                        6                        8                        6 
##          ENSG00000134874          ENSG00000134955          ENSG00000135046 
##                        2                        5                        6 
##          ENSG00000135074          ENSG00000135111          ENSG00000135114 
##                        4                        8                        4 
##          ENSG00000135245          ENSG00000135318          ENSG00000135406 
##                        5                        1                        4 
##          ENSG00000135437          ENSG00000135454          ENSG00000135547 
##                        2                        8                        6 
##          ENSG00000135622          ENSG00000135625          ENSG00000135636 
##                        2                        6                        5 
##          ENSG00000135919          ENSG00000135929          ENSG00000136040 
##                        7                        1                        7 
##          ENSG00000136144          ENSG00000136305          ENSG00000136449 
##                        2                        2                        6 
##          ENSG00000136770          ENSG00000136810          ENSG00000136826 
##                        2                        8                        2 
##          ENSG00000136828          ENSG00000136870          ENSG00000137070 
##                        8                        5                        2 
##          ENSG00000137078          ENSG00000137103          ENSG00000137135 
##                        5                        2                        7 
##          ENSG00000137142          ENSG00000137204          ENSG00000137216 
##                        4                        5                        8 
##          ENSG00000137285          ENSG00000137331          ENSG00000137393 
##                        8                        8                        5 
##          ENSG00000137460          ENSG00000137491          ENSG00000137501 
##                        4                        1                        4 
##          ENSG00000137547          ENSG00000137628          ENSG00000137802 
##                        1                        6                        2 
##          ENSG00000138074          ENSG00000138075          ENSG00000138080 
##                        1                        1                        1 
##          ENSG00000138111          ENSG00000138166          ENSG00000138336 
##                        8                        7                        5 
##          ENSG00000138363          ENSG00000138442          ENSG00000138449 
##                        8                        1                        8 
##          ENSG00000138496          ENSG00000138604          ENSG00000138615 
##                        2                        8                        4 
##          ENSG00000138678          ENSG00000138709          ENSG00000138778 
##                        2                        8                        7 
##          ENSG00000138821          ENSG00000138823          ENSG00000138834 
##                        4                        1                        8 
##          ENSG00000139055          ENSG00000139173          ENSG00000139182 
##                        7                        7                        8 
##          ENSG00000139219          ENSG00000139233          ENSG00000139269 
##                        7                        2                        1 
##          ENSG00000139292          ENSG00000139364          ENSG00000139428 
##                        5                        6                        8 
##          ENSG00000139514          ENSG00000139531          ENSG00000139540 
##                        1                        2                        1 
##          ENSG00000139880          ENSG00000139914          ENSG00000139926 
##                        4                        5                        2 
##          ENSG00000139988          ENSG00000140043          ENSG00000140092 
##                        2                        2                        8 
##          ENSG00000140107          ENSG00000140287          ENSG00000140323 
##                        1                        6                        7 
##          ENSG00000140398          ENSG00000140406          ENSG00000140807 
##                        5                        1                        8 
##          ENSG00000140836          ENSG00000141179          ENSG00000141448 
##                        7                        2                        2 
##          ENSG00000141485          ENSG00000141540          ENSG00000141542 
##                        8                        4                        2 
##          ENSG00000141574          ENSG00000141577          ENSG00000141699 
##                        5                        8                        8 
##          ENSG00000141736          ENSG00000141753          ENSG00000141873 
##                        1                        2                        8 
##          ENSG00000141934          ENSG00000141959          ENSG00000141994 
##                        4                        8                        1 
##          ENSG00000142182          ENSG00000142235          ENSG00000142273 
##                        4                        2                        2 
##          ENSG00000142449          ENSG00000142494          ENSG00000142552 
##                        5                        1                        3 
##          ENSG00000142661          ENSG00000142669          ENSG00000142675 
##                        8                        2                        2 
##          ENSG00000142871          ENSG00000143179          ENSG00000143222 
##                        8                        1                        8 
##          ENSG00000143318          ENSG00000143365          ENSG00000143369 
##                        6                        5                        4 
##          ENSG00000143375          ENSG00000143416          ENSG00000143418 
##                        2                        2                        1 
##          ENSG00000143434          ENSG00000143436          ENSG00000143554 
##                        8                        5                        5 
##          ENSG00000143603          ENSG00000143627          ENSG00000143819 
##                        6                        8                        8 
##          ENSG00000143845          ENSG00000143869          ENSG00000143921 
##                        8                        1                        5 
##          ENSG00000144035          ENSG00000144040          ENSG00000144285 
##                        5                        8                        5 
##          ENSG00000144381          ENSG00000144596          ENSG00000144647 
##                        8                        4                        8 
##          ENSG00000144730          ENSG00000144736          ENSG00000144749 
##                        4                        8                        1 
##          ENSG00000144815          ENSG00000144837          ENSG00000144857 
##                        2                        6                        4 
##          ENSG00000144893          ENSG00000145107          ENSG00000145217 
##                        4                        5                        7 
##          ENSG00000145391          ENSG00000145491          ENSG00000145506 
##                        4                        6                        2 
##          ENSG00000145632          ENSG00000145675          ENSG00000145757 
##                        2                        8                        3 
##          ENSG00000145860          ENSG00000145920          ENSG00000145949 
##                        5                        7                        3 
##          ENSG00000146001          ENSG00000146072          ENSG00000146166 
##                        3                        2                        4 
##          ENSG00000146216          ENSG00000146242          ENSG00000146416 
##                        5                        4                        8 
##          ENSG00000146592          ENSG00000146700          ENSG00000146755 
##                        4                        8                        7 
##          ENSG00000146828          ENSG00000146904          ENSG00000146955 
##                        8                        8                        6 
##          ENSG00000146966          ENSG00000147155          ENSG00000147206 
##                        6                        1                        6 
##          ENSG00000147255          ENSG00000147400          ENSG00000147465 
##                        1                        2                        5 
##          ENSG00000147676          ENSG00000147804          ENSG00000148218 
##                        8                        1                        8 
##          ENSG00000148225          ENSG00000148291          ENSG00000148468 
##                        2                        2                        1 
##          ENSG00000148484          ENSG00000148672          ENSG00000148677 
##                        8                        8                        2 
##          ENSG00000148730          ENSG00000148773          ENSG00000148795 
##                        8                        7                        8 
##          ENSG00000148832          ENSG00000148965          ENSG00000149043 
##                        2                        7                        5 
##          ENSG00000149212          ENSG00000149328          ENSG00000149451 
##                        7                        2                        4 
##          ENSG00000149476          ENSG00000149489          ENSG00000149503 
##                        1                        2                        8 
##          ENSG00000149564          ENSG00000149596          ENSG00000149599 
##                        2                        3                        4 
##          ENSG00000149742          ENSG00000149809          ENSG00000149926 
##                        5                        8                        6 
##          ENSG00000150510          ENSG00000150540          ENSG00000150867 
##                        4                        1                        8 
##          ENSG00000151117          ENSG00000151131          ENSG00000151320 
##                        4                        5                        4 
##          ENSG00000151348          ENSG00000151632          ENSG00000151690 
##                        8                        8                        7 
##          ENSG00000152076          ENSG00000152078          ENSG00000152082 
##                        4                        8                        5 
##          ENSG00000152192          ENSG00000152556          ENSG00000152763 
##                        2                        2                        4 
##          ENSG00000152778          ENSG00000152782          ENSG00000153012 
##                        4                        1                        6 
##          ENSG00000153060          ENSG00000153162          ENSG00000153233 
##                        6                        7                        6 
##          ENSG00000153363          ENSG00000153406          ENSG00000153714 
##                        4                        4                        5 
##          ENSG00000153885          ENSG00000153902          ENSG00000154153 
##                        5                        3                        4 
##          ENSG00000154217          ENSG00000154269          ENSG00000154319 
##                        2                        8                        6 
##          ENSG00000154553          ENSG00000154556          ENSG00000154856 
##                        2                        5                        8 
##          ENSG00000154864          ENSG00000154930          ENSG00000155016 
##                        5                        7                        4 
##          ENSG00000155115          ENSG00000155465          ENSG00000155629 
##                        5                        4                        8 
##          ENSG00000155846          ENSG00000155858          ENSG00000155893 
##                        4                        1                        2 
##          ENSG00000156222          ENSG00000156384          ENSG00000156413 
##                        5                        3                        5 
##          ENSG00000156453          ENSG00000156515          ENSG00000156966 
##                        2                        4                        6 
##          ENSG00000156973          ENSG00000157087          ENSG00000157193 
##                        2                        5                        5 
##          ENSG00000157303          ENSG00000157343          ENSG00000157510 
##                        2                        6                        4 
##          ENSG00000157514          ENSG00000157601          ENSG00000157625 
##                        4                        4                        8 
##          ENSG00000157657          ENSG00000157833          ENSG00000157856 
##                        4                        4                        6 
##          ENSG00000157873          ENSG00000158008          ENSG00000158106 
##                        8                        2                        2 
##          ENSG00000158158          ENSG00000158246          ENSG00000158296 
##                        8                        7                        8 
##          ENSG00000158373          ENSG00000158428          ENSG00000158458 
##                        8                        4                        6 
##          ENSG00000158560          ENSG00000158715          ENSG00000158813 
##                        6                        1                        2 
##          ENSG00000158955          ENSG00000159261          ENSG00000159403 
##                        4                        4                        4 
##          ENSG00000159409          ENSG00000159640          ENSG00000159650 
##                        4                        4                        5 
##          ENSG00000159733          ENSG00000159899          ENSG00000160211 
##                        4                        3                        8 
##          ENSG00000160255          ENSG00000160285          ENSG00000160323 
##                        8                        8                        2 
##          ENSG00000160446          ENSG00000160539          ENSG00000160602 
##                        2                        2                        5 
##          ENSG00000160688          ENSG00000160746          ENSG00000160870 
##                        8                        2                        7 
##          ENSG00000161011          ENSG00000161031          ENSG00000161082 
##                        8                        8                        4 
##          ENSG00000161180          ENSG00000161249          ENSG00000161328 
##                        4                        2                        2 
##          ENSG00000161513          ENSG00000161533          ENSG00000161638 
##                        8                        1                        4 
##          ENSG00000161682          ENSG00000162068          ENSG00000162241 
##                        8                        3                        2 
##          ENSG00000162267          ENSG00000162383          ENSG00000162391 
##                        8                        2                        7 
##          ENSG00000162413          ENSG00000162460          ENSG00000162461 
##                        1                        2                        4 
##          ENSG00000162482          ENSG00000162496          ENSG00000162551 
##                        4                        1                        2 
##          ENSG00000162572          ENSG00000162576          ENSG00000162599 
##                        8                        8                        7 
##          ENSG00000162676          ENSG00000162746          ENSG00000162772 
##                        8                        4                        8 
##          ENSG00000162783          ENSG00000162817          ENSG00000162869 
##                        8                        8                        2 
##          ENSG00000162889          ENSG00000162927          ENSG00000162959 
##                        8                        7                        1 
##          ENSG00000162976          ENSG00000163001          ENSG00000163002 
##                        7                        2                        8 
##          ENSG00000163040          ENSG00000163121          ENSG00000163235 
##                        6                        4                        1 
##          ENSG00000163295          ENSG00000163346          ENSG00000163431 
##                        4                        8                        6 
##          ENSG00000163531          ENSG00000163536          ENSG00000163577 
##                        4                        8                        4 
##          ENSG00000163586          ENSG00000163624          ENSG00000163631 
##                        7                        2                        8 
##          ENSG00000163666          ENSG00000163701          ENSG00000163792 
##                        6                        5                        5 
##          ENSG00000163884          ENSG00000163959          ENSG00000163964 
##                        2                        7                        2 
##          ENSG00000164007          ENSG00000164039          ENSG00000164050 
##                        8                        2                        1 
##          ENSG00000164078          ENSG00000164105          ENSG00000164125 
##                        2                        5                        1 
##          ENSG00000164236          ENSG00000164266          ENSG00000164287 
##                        4                        4                        6 
##          ENSG00000164379          ENSG00000164403          ENSG00000164406 
##                        8                        8                        7 
##          ENSG00000164484          ENSG00000164638          ENSG00000164674 
##                        4                        8                        4 
##          ENSG00000164687          ENSG00000164841          ENSG00000164949 
##                        2                        6                        4 
##          ENSG00000165029          ENSG00000165092          ENSG00000165097 
##                        1                        1                        2 
##          ENSG00000165105          ENSG00000165140          ENSG00000165169 
##                        6                        4                        4 
##          ENSG00000165215          ENSG00000165269          ENSG00000165272 
##                        2                        2                        7 
##          ENSG00000165283          ENSG00000165434          ENSG00000165474 
##                        1                        7                        6 
##          ENSG00000165478          ENSG00000165526          ENSG00000165548 
##                        8                        8                        2 
##          ENSG00000165630          ENSG00000165632          ENSG00000165695 
##                        2                        8                        4 
##          ENSG00000165732          ENSG00000165914          ENSG00000165929 
##                        8                        2                        6 
##          ENSG00000165949          ENSG00000166035          ENSG00000166046 
##                        7                        5                        8 
##          ENSG00000166126          ENSG00000166128          ENSG00000166145 
##                        1                        2                        2 
##          ENSG00000166272          ENSG00000166278          ENSG00000166289 
##                        8                        8                        2 
##          ENSG00000166311          ENSG00000166313          ENSG00000166415 
##                        8                        4                        5 
##          ENSG00000166508          ENSG00000166532          ENSG00000166589 
##                        1                        4                        8 
##          ENSG00000166592          ENSG00000166780          ENSG00000166801 
##                        4                        2                        5 
##          ENSG00000166816          ENSG00000166823          ENSG00000166866 
##                        2                        6                        8 
##          ENSG00000166963          ENSG00000167065          ENSG00000167123 
##                        4                        4                        2 
##          ENSG00000167131          ENSG00000167183          ENSG00000167311 
##                        4                        5                        4 
##          ENSG00000167371          ENSG00000167380          ENSG00000167508 
##                        4                        5                        8 
##          ENSG00000167535          ENSG00000167552          ENSG00000167588 
##                        4                        7                        5 
##          ENSG00000167608          ENSG00000167632          ENSG00000167664 
##                        3                        2                        8 
##          ENSG00000167740          ENSG00000167780          ENSG00000167964 
##                        2                        5                        7 
##          ENSG00000167996          ENSG00000168005          ENSG00000168028 
##                        8                        8                        1 
##          ENSG00000168060          ENSG00000168077          ENSG00000168081 
##                        5                        5                        6 
##          ENSG00000168461          ENSG00000168491          ENSG00000168497 
##                        7                        4                        5 
##          ENSG00000168502          ENSG00000168505          ENSG00000168646 
##                        7                        6                        8 
##          ENSG00000168675          ENSG00000168778          ENSG00000168785 
##                        5                        2                        4 
##          ENSG00000168916          ENSG00000168938          ENSG00000169026 
##                        6                        2                        3 
##          ENSG00000169174          ENSG00000169213          ENSG00000169231 
##                        1                        7                        4 
##          ENSG00000169291          ENSG00000169379          ENSG00000169403 
##                        6                        2                        8 
##          ENSG00000169548          ENSG00000169562          ENSG00000169583 
##                        4                        1                        2 
##          ENSG00000169688          ENSG00000169692          ENSG00000169856 
##                        4                        8                        5 
##          ENSG00000169884          ENSG00000169891          ENSG00000169962 
##                        6                        4                        5 
##          ENSG00000169994          ENSG00000170004          ENSG00000170074 
##                        4                        8                        3 
##          ENSG00000170099          ENSG00000170153          ENSG00000170271 
##                        8                        6                        5 
##          ENSG00000170482          ENSG00000170485          ENSG00000170522 
##                        5                        5                        8 
##          ENSG00000170542          ENSG00000170629          ENSG00000170667 
##                        8                        4                        2 
##          ENSG00000170835          ENSG00000170965          ENSG00000171004 
##                        5                        4                        7 
##          ENSG00000171016          ENSG00000171219          ENSG00000171223 
##                        6                        8                        7 
##          ENSG00000171241          ENSG00000171314          ENSG00000171357 
##                        7                        8                        6 
##          ENSG00000171385          ENSG00000171425          ENSG00000171435 
##                        6                        1                        4 
##          ENSG00000171450          ENSG00000171462          ENSG00000171557 
##                        4                        5                        8 
##          ENSG00000171560          ENSG00000171564          ENSG00000171621 
##                        8                        8                        8 
##          ENSG00000171703          ENSG00000171766          ENSG00000171798 
##                        2                        5                        4 
##          ENSG00000171813          ENSG00000171840          ENSG00000171847 
##                        1                        3                        6 
##          ENSG00000171863          ENSG00000172031          ENSG00000172236 
##                        8                        6                        6 
##          ENSG00000172296          ENSG00000172331          ENSG00000172409 
##                        1                        4                        2 
##          ENSG00000172478          ENSG00000172482          ENSG00000172638 
##                        5                        8                        8 
##          ENSG00000172757          ENSG00000172794          ENSG00000172818 
##                        8                        2                        6 
##          ENSG00000172824          ENSG00000172889          ENSG00000172890 
##                        4                        2                        8 
##          ENSG00000172915          ENSG00000172955          ENSG00000173065 
##                        2                        5                        2 
##          ENSG00000173110          ENSG00000173156          ENSG00000173210 
##                        7                        2                        4 
##          ENSG00000173212          ENSG00000173227          ENSG00000173264 
##                        5                        4                        8 
##          ENSG00000173272          ENSG00000173320          ENSG00000173457 
##                        5                        6                        5 
##          ENSG00000173567          ENSG00000173597          ENSG00000173714 
##                        2                        7                        4 
##          ENSG00000173762          ENSG00000173805          ENSG00000173825 
##                        2                        4                        2 
##          ENSG00000173930          ENSG00000174004          ENSG00000174125 
##                        2                        6                        6 
##          ENSG00000174130          ENSG00000174460          ENSG00000174672 
##                        6                        4                        2 
##          ENSG00000174744          ENSG00000174827          ENSG00000174917 
##                        8                        5                        2 
##          ENSG00000174938          ENSG00000174939          ENSG00000175063 
##                        7                        2                        5 
##          ENSG00000175084          ENSG00000175106          ENSG00000175155 
##                        4                        5                        8 
##          ENSG00000175170          ENSG00000175197          ENSG00000175279 
##                        4                        7                        2 
##          ENSG00000175287          ENSG00000175294          ENSG00000175318 
##                        5                        6                        4 
##          ENSG00000175329          ENSG00000175336          ENSG00000175416 
##                        5                        7                        2 
##          ENSG00000175550          ENSG00000175575          ENSG00000175592 
##                        2                        1                        5 
##          ENSG00000175938          ENSG00000175946          ENSG00000176046 
##                        5                        6                        7 
##          ENSG00000176058          ENSG00000176134          ENSG00000176153 
##                        8                        7                        7 
##          ENSG00000176473          ENSG00000176593          ENSG00000176641 
##                        4                        3                        6 
##          ENSG00000176658          ENSG00000176728          ENSG00000176834 
##                        8                        6                        8 
##          ENSG00000176845          ENSG00000176945          ENSG00000176978 
##                        4                        2                        5 
##          ENSG00000177000          ENSG00000177076          ENSG00000177238 
##                        8                        2                        6 
##          ENSG00000177283          ENSG00000177570          ENSG00000177595 
##                        4                        6                        1 
##          ENSG00000177606          ENSG00000177679          ENSG00000177685 
##                        8                        2                        8 
##          ENSG00000177700          ENSG00000177706          ENSG00000177752 
##                        8                        5                        6 
##          ENSG00000177943          ENSG00000177981          ENSG00000177984 
##                        1                        2                        7 
##          ENSG00000178150          ENSG00000178301          ENSG00000178531 
##                        2                        2                        2 
##          ENSG00000178597          ENSG00000178695          ENSG00000178718 
##                        4                        6                        4 
##          ENSG00000178977          ENSG00000178996          ENSG00000179044 
##                        4                        2                        4 
##          ENSG00000179091          ENSG00000179148          ENSG00000179178 
##                        8                        4                        4 
##          ENSG00000179240          ENSG00000179292          ENSG00000179314 
##                        2                        2                        4 
##          ENSG00000179344          ENSG00000179388          ENSG00000179520 
##                        4                        6                        4 
##          ENSG00000179598          ENSG00000179776          ENSG00000179913 
##                        5                        4                        5 
##          ENSG00000180044          ENSG00000180096          ENSG00000180155 
##                        6                        2                        2 
##          ENSG00000180176          ENSG00000180198          ENSG00000180264 
##                        6                        1                        6 
##          ENSG00000180425          ENSG00000180432          ENSG00000180448 
##                        2                        3                        1 
##          ENSG00000180694          ENSG00000180767          ENSG00000181019 
##                        1                        1                        1 
##          ENSG00000181029          ENSG00000181392          ENSG00000181408 
##                        8                        2                        3 
##          ENSG00000181472          ENSG00000181577          ENSG00000181649 
##                        5                        8                        2 
##          ENSG00000181856          ENSG00000182118          ENSG00000182156 
##                        2                        6                        5 
##          ENSG00000182162 ENSG00000182162.11_PAR_Y          ENSG00000182240 
##                        5                        5                        6 
##          ENSG00000182253          ENSG00000182327          ENSG00000182379 
##                        2                        1                        2 
##          ENSG00000182459          ENSG00000182472          ENSG00000182492 
##                        2                        7                        6 
##          ENSG00000182557          ENSG00000182782          ENSG00000182795 
##                        6                        3                        2 
##          ENSG00000182796          ENSG00000182809          ENSG00000182912 
##                        2                        2                        2 
##          ENSG00000183011          ENSG00000183019          ENSG00000183077 
##                        2                        6                        1 
##          ENSG00000183128          ENSG00000183153          ENSG00000183307 
##                        6                        7                        6 
##          ENSG00000183401          ENSG00000183421          ENSG00000183496 
##                        2                        8                        2 
##          ENSG00000183508          ENSG00000183549          ENSG00000183598 
##                        4                        5                        5 
##          ENSG00000183629          ENSG00000183684          ENSG00000183688 
##                        6                        5                        8 
##          ENSG00000183747          ENSG00000183779          ENSG00000183889 
##                        4                        8                        1 
##          ENSG00000183914          ENSG00000183971          ENSG00000184160 
##                        6                        2                        2 
##          ENSG00000184194          ENSG00000184226          ENSG00000184227 
##                        4                        4                        1 
##          ENSG00000184232          ENSG00000184270          ENSG00000184271 
##                        8                        5                        6 
##          ENSG00000184357          ENSG00000184371          ENSG00000184384 
##                        5                        2                        5 
##          ENSG00000184470          ENSG00000184489          ENSG00000184502 
##                        5                        5                        4 
##          ENSG00000184602          ENSG00000184678          ENSG00000184697 
##                        7                        7                        8 
##          ENSG00000184897          ENSG00000184949          ENSG00000184985 
##                        8                        4                        5 
##          ENSG00000184986          ENSG00000185022          ENSG00000185085 
##                        2                        8                        2 
##          ENSG00000185100          ENSG00000185112          ENSG00000185115 
##                        5                        4                        2 
##          ENSG00000185130          ENSG00000185187          ENSG00000185189 
##                        5                        3                        2 
##          ENSG00000185201          ENSG00000185262          ENSG00000185303 
##                        5                        5                        6 
##          ENSG00000185432          ENSG00000185483          ENSG00000185532 
##                        1                        4                        6 
##          ENSG00000185565          ENSG00000185633          ENSG00000185686 
##                        3                        5                        4 
##          ENSG00000185813          ENSG00000185818          ENSG00000185875 
##                        1                        2                        2 
##          ENSG00000185880          ENSG00000185885          ENSG00000185909 
##                        2                        5                        2 
##          ENSG00000185986          ENSG00000186047          ENSG00000186106 
##                        5                        3                        8 
##          ENSG00000186115          ENSG00000186204          ENSG00000186212 
##                        7                        8                        4 
##          ENSG00000186326          ENSG00000186529          ENSG00000186564 
##                        4                        5                        4 
##          ENSG00000186567          ENSG00000186577          ENSG00000186854 
##                        3                        2                        3 
##          ENSG00000186868          ENSG00000186897          ENSG00000186910 
##                        7                        4                        5 
##          ENSG00000186918          ENSG00000186994          ENSG00000186998 
##                        8                        4                        4 
##          ENSG00000187045          ENSG00000187097          ENSG00000187134 
##                        5                        8                        8 
##          ENSG00000187189          ENSG00000187231          ENSG00000187240 
##                        2                        8                        4 
##          ENSG00000187244          ENSG00000187391          ENSG00000187475 
##                        8                        5                        6 
##          ENSG00000187479          ENSG00000187550          ENSG00000187608 
##                        7                        2                        8 
##          ENSG00000187627          ENSG00000187650          ENSG00000187678 
##                        7                        2                        8 
##          ENSG00000187686          ENSG00000187720          ENSG00000187800 
##                        4                        4                        4 
##          ENSG00000187867          ENSG00000187908          ENSG00000187955 
##                        5                        6                        6 
##          ENSG00000187994          ENSG00000187997          ENSG00000188064 
##                        5                        3                        6 
##          ENSG00000188107          ENSG00000188338          ENSG00000188372 
##                        3                        8                        3 
##          ENSG00000188385          ENSG00000188582          ENSG00000188641 
##                        4                        1                        2 
##          ENSG00000188760          ENSG00000188818          ENSG00000188827 
##                        8                        4                        2 
##          ENSG00000188833          ENSG00000188878          ENSG00000189120 
##                        7                        2                        4 
##          ENSG00000189143          ENSG00000189159          ENSG00000189221 
##                        8                        8                        1 
##          ENSG00000196136          ENSG00000196167          ENSG00000196177 
##                        7                        3                        1 
##          ENSG00000196208          ENSG00000196220          ENSG00000196337 
##                        1                        4                        4 
##          ENSG00000196415          ENSG00000196456          ENSG00000196497 
##                        6                        2                        8 
##          ENSG00000196544          ENSG00000196611          ENSG00000196660 
##                        2                        6                        1 
##          ENSG00000196670          ENSG00000196693          ENSG00000196747 
##                        2                        5                        5 
##          ENSG00000196814          ENSG00000196839          ENSG00000196866 
##                        1                        2                        5 
##          ENSG00000196872          ENSG00000196876          ENSG00000196917 
##                        2                        4                        5 
##          ENSG00000196967          ENSG00000196972          ENSG00000196990 
##                        2                        4                        6 
##          ENSG00000197063          ENSG00000197085          ENSG00000197253 
##                        8                        3                        4 
##          ENSG00000197291          ENSG00000197296          ENSG00000197301 
##                        3                        8                        3 
##          ENSG00000197479          ENSG00000197557          ENSG00000197568 
##                        4                        4                        4 
##          ENSG00000197653          ENSG00000197766          ENSG00000197858 
##                        4                        2                        8 
##          ENSG00000198074          ENSG00000198075          ENSG00000198203 
##                        7                        5                        5 
##          ENSG00000198208          ENSG00000198221          ENSG00000198431 
##                        2                        2                        8 
##          ENSG00000198576          ENSG00000198624          ENSG00000198648 
##                        2                        8                        5 
##          ENSG00000198663          ENSG00000198691          ENSG00000198695 
##                        8                        4                        7 
##          ENSG00000198756          ENSG00000198780          ENSG00000198805 
##                        5                        7                        2 
##          ENSG00000198832          ENSG00000198848          ENSG00000198892 
##                        7                        7                        4 
##          ENSG00000198910          ENSG00000198911          ENSG00000198932 
##                        8                        8                        3 
##          ENSG00000198948          ENSG00000198951          ENSG00000198963 
##                        4                        2                        6 
##          ENSG00000203690          ENSG00000203709          ENSG00000203722 
##                        4                        5                        6 
##          ENSG00000203739          ENSG00000203814          ENSG00000203852 
##                        3                        5                        7 
##          ENSG00000203867          ENSG00000204044          ENSG00000204099 
##                        4                        4                        1 
##          ENSG00000204103          ENSG00000204113          ENSG00000204128 
##                        4                        3                        1 
##          ENSG00000204176          ENSG00000204177          ENSG00000204195 
##                        3                        5                        6 
##          ENSG00000204228          ENSG00000204248          ENSG00000204257 
##                        2                        4                        4 
##          ENSG00000204388          ENSG00000204389          ENSG00000204444 
##                        8                        8                        8 
##          ENSG00000204525          ENSG00000204681          ENSG00000204682 
##                        8                        4                        5 
##          ENSG00000204852          ENSG00000204859          ENSG00000204869 
##                        2                        2                        4 
##          ENSG00000204882          ENSG00000204956          ENSG00000204967 
##                        4                        4                        4 
##          ENSG00000204991          ENSG00000205116          ENSG00000205181 
##                        8                        8                        4 
##          ENSG00000205269          ENSG00000205277          ENSG00000205436 
##                        8                        4                        8 
##          ENSG00000205559          ENSG00000205639          ENSG00000205710 
##                        2                        3                        3 
##          ENSG00000205746          ENSG00000205795          ENSG00000205808 
##                        1                        4                        2 
##          ENSG00000205918          ENSG00000205922          ENSG00000205978 
##                        5                        6                        2 
##          ENSG00000206077          ENSG00000206127          ENSG00000206172 
##                        4                        7                        7 
##          ENSG00000206561          ENSG00000212724          ENSG00000213339 
##                        6                        6                        1 
##          ENSG00000213366          ENSG00000213398          ENSG00000213420 
##                        2                        5                        2 
##          ENSG00000213549          ENSG00000213563          ENSG00000213762 
##                        6                        8                        2 
##          ENSG00000213859          ENSG00000213889          ENSG00000213930 
##                        8                        2                        8 
##          ENSG00000213939          ENSG00000213949          ENSG00000213999 
##                        6                        5                        2 
##          ENSG00000214212          ENSG00000214279          ENSG00000214313 
##                        6                        2                        4 
##          ENSG00000214357          ENSG00000214402          ENSG00000214491 
##                        4                        6                        5 
##          ENSG00000214706          ENSG00000215105          ENSG00000215906 
##                        1                        4                        5 
##          ENSG00000215910          ENSG00000218510          ENSG00000219607 
##                        4                        2                        5 
##          ENSG00000220323          ENSG00000220412          ENSG00000220785 
##                        7                        6                        3 
##          ENSG00000221869          ENSG00000221890          ENSG00000221990 
##                        5                        8                        2 
##          ENSG00000222009          ENSG00000223612          ENSG00000223638 
##                        5                        6                        6 
##          ENSG00000223745          ENSG00000224078          ENSG00000224177 
##                        2                        8                        6 
##          ENSG00000224376          ENSG00000224397          ENSG00000224430 
##                        6                        4                        3 
##          ENSG00000224596          ENSG00000224616          ENSG00000224914 
##                        2                        4                        2 
##          ENSG00000225026          ENSG00000225138          ENSG00000225431 
##                        6                        2                        6 
##          ENSG00000225595          ENSG00000225630          ENSG00000225756 
##                        6                        4                        5 
##          ENSG00000225793          ENSG00000226015          ENSG00000226043 
##                        4                        3                        6 
##          ENSG00000226137          ENSG00000226510          ENSG00000226555 
##                        2                        4                        3 
##          ENSG00000226742          ENSG00000226976          ENSG00000227825 
##                        3                        6                        4 
##          ENSG00000227827          ENSG00000228146          ENSG00000228412 
##                        1                        3                        4 
##          ENSG00000228451          ENSG00000228549          ENSG00000228594 
##                        2                        4                        2 
##          ENSG00000228695          ENSG00000228697          ENSG00000228705 
##                        6                        3                        4 
##          ENSG00000228804          ENSG00000228903          ENSG00000229356 
##                        3                        8                        6 
##          ENSG00000229474          ENSG00000229666          ENSG00000230091 
##                        6                        6                        6 
##          ENSG00000230359          ENSG00000230590          ENSG00000230882 
##                        4                        7                        5 
##          ENSG00000231066          ENSG00000231305          ENSG00000231515 
##                        6                        3                        6 
##          ENSG00000231683          ENSG00000231690          ENSG00000231969 
##                        7                        2                        3 
##          ENSG00000232453          ENSG00000232624          ENSG00000232767 
##                        6                        4                        4 
##          ENSG00000233058          ENSG00000233392          ENSG00000233421 
##                        2                        5                        4 
##          ENSG00000233695          ENSG00000233922          ENSG00000233990 
##                        4                        4                        3 
##          ENSG00000234199          ENSG00000234438          ENSG00000234473 
##                        4                        6                        4 
##          ENSG00000234899          ENSG00000234996          ENSG00000235092 
##                        4                        2                        7 
##          ENSG00000235098          ENSG00000235142          ENSG00000235257 
##                        5                        3                        3 
##          ENSG00000235390          ENSG00000235568          ENSG00000235890 
##                        4                        6                        2 
##          ENSG00000235899          ENSG00000236107          ENSG00000236279 
##                        4                        7                        6 
##          ENSG00000236384          ENSG00000236714          ENSG00000237187 
##                        4                        3                        5 
##          ENSG00000237289          ENSG00000237361          ENSG00000237390 
##                        4                        3                        3 
##          ENSG00000237471          ENSG00000237522          ENSG00000237595 
##                        4                        3                        4 
##          ENSG00000237643          ENSG00000237686          ENSG00000237693 
##                        4                        2                        4 
##          ENSG00000237989          ENSG00000238062          ENSG00000238164 
##                        6                        6                        7 
##          ENSG00000238178          ENSG00000239282          ENSG00000239713 
##                        6                        4                        4 
##          ENSG00000239779          ENSG00000240021          ENSG00000240303 
##                        8                        6                        1 
##          ENSG00000241388          ENSG00000241635          ENSG00000241697 
##                        5                        7                        4 
##          ENSG00000241764          ENSG00000242071          ENSG00000242173 
##                        4                        7                        6 
##          ENSG00000242220          ENSG00000242242          ENSG00000243317 
##                        3                        4                        2 
##          ENSG00000243359          ENSG00000243649          ENSG00000243818 
##                        6                        8                        3 
##          ENSG00000243955          ENSG00000244062          ENSG00000244198 
##                        7                        8                        3 
##          ENSG00000244242          ENSG00000244411          ENSG00000244509 
##                        4                        5                        8 
##          ENSG00000244968          ENSG00000246526          ENSG00000246705 
##                        4                        4                        2 
##          ENSG00000246790          ENSG00000246877          ENSG00000247240 
##                        3                        6                        4 
##          ENSG00000247735          ENSG00000248092          ENSG00000248429 
##                        4                        5                        7 
##          ENSG00000248485          ENSG00000249201          ENSG00000249715 
##                        2                        5                        4 
##          ENSG00000249790          ENSG00000249992          ENSG00000250318 
##                        2                        6                        6 
##          ENSG00000250799          ENSG00000250903          ENSG00000251034 
##                        1                        5                        3 
##          ENSG00000251165          ENSG00000251201          ENSG00000251381 
##                        3                        5                        4 
##          ENSG00000251493          ENSG00000253210          ENSG00000253313 
##                        4                        4                        2 
##          ENSG00000253516          ENSG00000253953          ENSG00000254042 
##                        3                        2                        5 
##          ENSG00000254054          ENSG00000254109          ENSG00000254192 
##                        7                        4                        4 
##          ENSG00000254224          ENSG00000254635          ENSG00000254681 
##                        3                        2                        1 
##          ENSG00000254990          ENSG00000255121          ENSG00000255198 
##                        3                        4                        6 
##          ENSG00000255398          ENSG00000255498          ENSG00000255959 
##                        5                        3                        3 
##          ENSG00000256124          ENSG00000256204          ENSG00000256249 
##                        4                        6                        3 
##          ENSG00000256340          ENSG00000257017          ENSG00000257093 
##                        8                        7                        5 
##          ENSG00000257335          ENSG00000257433          ENSG00000257512 
##                        7                        4                        2 
##          ENSG00000257595          ENSG00000257702          ENSG00000258451 
##                        6                        2                        3 
##          ENSG00000258655          ENSG00000258702          ENSG00000258947 
##                        4                        6                        8 
##          ENSG00000259352          ENSG00000259370          ENSG00000259687 
##                        3                        3                        6 
##          ENSG00000259877          ENSG00000259884          ENSG00000259974 
##                        4                        3                        7 
##          ENSG00000260022          ENSG00000260081          ENSG00000260252 
##                        3                        4                        6 
##          ENSG00000260293          ENSG00000260475          ENSG00000260563 
##                        5                        6                        2 
##          ENSG00000260573          ENSG00000260630          ENSG00000260708 
##                        5                        2                        4 
##          ENSG00000260777          ENSG00000260778          ENSG00000260804 
##                        6                        6                        5 
##          ENSG00000260852          ENSG00000261071          ENSG00000261150 
##                        5                        4                        7 
##          ENSG00000261189          ENSG00000261242          ENSG00000261644 
##                        7                        6                        6 
##          ENSG00000261655          ENSG00000261668          ENSG00000261739 
##                        6                        3                        6 
##          ENSG00000261934          ENSG00000262155          ENSG00000263004 
##                        6                        4                        6 
##          ENSG00000263155          ENSG00000264207          ENSG00000264404 
##                        6                        4                        4 
##          ENSG00000264578          ENSG00000265185          ENSG00000265692 
##                        7                        2                        6 
##          ENSG00000265794          ENSG00000266378          ENSG00000266709 
##                        6                        4                        5 
##          ENSG00000266714          ENSG00000266964          ENSG00000266970 
##                        8                        5                        6 
##          ENSG00000267121          ENSG00000267199          ENSG00000267221 
##                        6                        3                        4 
##          ENSG00000267296          ENSG00000267374          ENSG00000267534 
##                        2                        7                        2 
##          ENSG00000267576          ENSG00000267934          ENSG00000268230 
##                        7                        3                        1 
##          ENSG00000268350          ENSG00000268926          ENSG00000270276 
##                        7                        4                        7 
##          ENSG00000270885          ENSG00000270964          ENSG00000271122 
##                        1                        4                        2 
##          ENSG00000271303          ENSG00000271614          ENSG00000271646 
##                        8                        2                        4 
##          ENSG00000271755          ENSG00000271780          ENSG00000271855 
##                        6                        4                        6 
##          ENSG00000272031          ENSG00000272398          ENSG00000272568 
##                        4                        7                        3 
##          ENSG00000272686          ENSG00000272692          ENSG00000272902 
##                        4                        4                        4 
##          ENSG00000273066          ENSG00000273253          ENSG00000273259 
##                        5                        3                        2 
##          ENSG00000273381          ENSG00000273416          ENSG00000273456 
##                        5                        6                        4 
##          ENSG00000273604          ENSG00000273702          ENSG00000273703 
##                        8                        4                        5 
##          ENSG00000273706          ENSG00000273888          ENSG00000274080 
##                        6                        6                        5 
##          ENSG00000274180          ENSG00000274317          ENSG00000274444 
##                        2                        3                        6 
##          ENSG00000274922          ENSG00000274979          ENSG00000275152 
##                        2                        5                        5 
##          ENSG00000275395          ENSG00000275401          ENSG00000275832 
##                        5                        6                        4 
##          ENSG00000276045          ENSG00000276067          ENSG00000276076 
##                        2                        6                        7 
##          ENSG00000276141          ENSG00000276231          ENSG00000276410 
##                        5                        6                        5 
##          ENSG00000276966          ENSG00000276980          ENSG00000277224 
##                        5                        5                        5 
##          ENSG00000277287          ENSG00000277352          ENSG00000277453 
##                        4                        5                        6 
##          ENSG00000277561          ENSG00000277578          ENSG00000278384 
##                        4                        6                        5 
##          ENSG00000278463          ENSG00000278535          ENSG00000278619 
##                        8                        8                        5 
##          ENSG00000278677          ENSG00000278964          ENSG00000278967 
##                        8                        4                        3 
##          ENSG00000279041          ENSG00000279879          ENSG00000279894 
##                        3                        6                        3 
##          ENSG00000279925          ENSG00000280132          ENSG00000280161 
##                        6                        3                        4 
##          ENSG00000280245          ENSG00000280383          ENSG00000280399 
##                        3                        5                        3 
##          ENSG00000281344          ENSG00000282034          ENSG00000282057 
##                        7                        5                        6 
##          ENSG00000282807          ENSG00000282965          ENSG00000283071 
##                        6                        4                        4 
##          ENSG00000283154          ENSG00000283183          ENSG00000283632 
##                        6                        6                        4 
##          ENSG00000283930          ENSG00000284308          ENSG00000284713 
##                        5                        4                        6 
##          ENSG00000284906          ENSG00000285230          ENSG00000285513 
##                        5                        2                        7 
##          ENSG00000285517          ENSG00000285730          ENSG00000286116 
##                        5                        4                        3 
##          ENSG00000286190          ENSG00000286214          ENSG00000286388 
##                        7                        3                        2 
##          ENSG00000286403          ENSG00000286522          ENSG00000286546 
##                        7                        7                        7 
##          ENSG00000286623          ENSG00000286724          ENSG00000286733 
##                        6                        3                        5 
##          ENSG00000286757          ENSG00000287001          ENSG00000287080 
##                        3                        5                        7 
##          ENSG00000287097          ENSG00000287134          ENSG00000287263 
##                        4                        3                        3 
##          ENSG00000287419          ENSG00000287426          ENSG00000287529 
##                        6                        2                        4 
##          ENSG00000287591          ENSG00000287979          ENSG00000288009 
##                        7                        4                        6 
##          ENSG00000288622 
##                        4 
## 
## $cluster.algorithm.used
## [1] "Mclust_8"
## 
## $groups
##           Control NaBut
## Nt1_1           1     0
## Nt1_2           1     0
## Nt1_3           1     0
## Nt2_1           1     0
## Nt2_2           1     0
## Nt2_3           1     0
## Nt3_1           1     0
## Nt3_2           1     0
## Nt3_3           1     0
## Nt10_1          1     0
## Nt10_2          1     0
## Nt10_3          1     0
## NaBut1_1        0     1
## NaBut1_2        0     1
## NaBut1_3        0     1
## NaBut2_1        0     1
## NaBut2_2        0     1
## NaBut2_3        0     1
## NaBut3_1        0     1
## NaBut3_2        0     1
## NaBut3_3        0     1
## NaBut10_1       0     1
## NaBut10_2       0     1
## NaBut10_3       0     1
```

NaBut vs Control - Genes that have different time dependent trajectories between conditions - 6230 genes

```
# number of significant
sigs$sig.genes$NaButvsControl$g
```

```
## [1] 6161
```

```
# plot these genes 
res <- see.genes(sigs$sig.genes$NaButvsControl, show.fit = T, dis =design$dis, cluster.method="Mclust", cluster.data = 1, k.mclust=TRUE)
```

```
# output
write.csv(sigs$sig.genes$NaButvsControl$sig.profiles, "/Users/ronaldcutler/Library/CloudStorage//My\ Drive/Sidoli_Lab/Projects/Chromatin\ Decondensation/Analysis/Gencode/time-series_NaBut_vs_control_sig.csv")

# heatmap of all sig genes 
dds_vst.mat.var <- sigs$sig.genes$NaButvsControl$sig.profiles

# determine optimal clusters
mclustBIC(dds_vst.mat.var)
```

```
## Bayesian Information Criterion (BIC): 
##         EII       VII       EEI       VEI       EVI       VVI       EEE
## 1 -618486.7 -618486.7 -617120.9 -617120.9 -617120.9 -617120.9 -111431.9
## 2 -474826.9 -437838.2 -474006.3 -432677.9 -473644.1 -416121.0 -109096.0
## 3 -416318.3 -368320.9 -408510.8 -356330.7 -406593.4 -342993.6 -108518.1
## 4 -385509.4 -331591.1 -368560.7 -309814.6 -365691.4        NA -108434.1
## 5 -365074.8 -313823.3 -340650.1 -278232.9 -349952.9        NA -104427.9
## 6 -343726.4 -291686.2 -321416.0 -255805.4 -322818.8        NA -102720.1
## 7 -326065.4 -271954.1 -318480.2 -239665.6 -310361.8        NA -102160.9
## 8 -316592.6 -259761.7 -297688.6 -227726.2 -294528.2        NA -101372.2
## 9 -295368.9 -250087.0 -290430.6 -217634.3        NA        NA -101100.6
##          VEE        EVE       VVE        EEV        VEV        EVV       VVV
## 1 -111431.86 -111431.86 -111431.9 -111431.86 -111431.86 -111431.86 -111431.9
## 2  -96670.97  -90541.29        NA  -93293.37  -63841.74  -86156.97        NA
## 3  -91764.54  -83637.82        NA  -87052.87  -51204.29  -79534.45        NA
## 4  -89024.08  -81166.49        NA  -85237.31  -46648.60  -76981.82        NA
## 5  -86940.07         NA        NA  -81637.63  -44243.78         NA        NA
## 6  -85066.18         NA        NA  -81585.34  -40370.99         NA        NA
## 7  -86587.96         NA        NA  -78159.40  -40225.43         NA        NA
## 8  -83852.14         NA        NA  -77911.27  -39657.50         NA        NA
## 9         NA         NA        NA  -77688.74  -37500.02         NA        NA
## 
## Top 3 models based on the BIC criterion: 
##     VEV,9     VEV,8     VEV,7 
## -37500.02 -39657.50 -40225.43
```

```
# kmeans clustering
set.seed(123)
clust <- kmeans(dds_vst.mat.var, 8)
old <- 1:8
new <- c("C","F","D","A","B","H","E","G")
clust$cluster[clust$cluster %in% old] <- new[match(clust$cluster, old, nomatch = 0)]
split <- factor(clust$cluster, levels=rev(c("A","B","C","D","E","F","G","H")))

# output with clusters
dds_vst.mat.var.clust <- dds_vst.mat.var
dds_vst.mat.var.clust$cluster <- split
dds_vst.mat.var.clust$symbol <- names[match(rownames(dds_vst.mat.var.clust), names$ensembl_gene_id), "external_gene_name"]
dds_vst.mat.var.clust$entrezgene_id <- names[match(rownames(dds_vst.mat.var.clust), names$ensembl_gene_id), "entrezgene_id"]
dds_vst.mat.var.clust$gene_biotype <- names[match(rownames(dds_vst.mat.var.clust), names$ensembl_gene_id), "gene_biotype"]
write.csv(dds_vst.mat.var.clust, "/Users/ronaldcutler/Library/CloudStorage//My\ Drive/Sidoli_Lab/Projects/Chromatin\ Decondensation/Analysis/Gencode/time-series_NaBut_vs_control_sig_cluster.csv")

# heatmap split up by cluster
library(circlize)
```

```
## ========================================
## circlize version 0.4.14
## CRAN page: https://cran.r-project.org/package=circlize
## Github page: https://github.com/jokergoo/circlize
## Documentation: https://jokergoo.github.io/circlize_book/book/
## 
## If you use it in published research, please cite:
## Gu, Z. circlize implements and enhances circular visualization
##   in R. Bioinformatics 2014.
## 
## This message can be suppressed by:
##   suppressPackageStartupMessages(library(circlize))
## ========================================
```

```
colnames(dds_vst.mat.var) <- sampleTable$sample
col_fun = brewer.pal(4, "Purples")
ha <- HeatmapAnnotation(Condition = sampleTable$condition, Time = sampleTable$time,
                        col = list(Condition = c("Nt" = "#7f7f7f", "NaBut" = "#006837"),
                                   Time = c("1" = "#F2F0F7", "2" = "#CBC9E2", "3" = "#9E9AC8", "10" = "#6A51A3")))
scaled.mat <- t(scale(t(dds_vst.mat.var),center=TRUE,scale=TRUE))
Heatmap(scaled.mat, 
        name = "Z-score", #title of legend
        row_names_gp = gpar(fontsize = 7), # Text size for row names
        split = split,
        column_split = factor(sampleTable$condition, levels = c("Nt", "NaBut")),
        cluster_row_slices = FALSE,
        top_annotation = ha,
        border = TRUE,
        cluster_rows = TRUE,
        cluster_columns = FALSE,
        show_row_names = FALSE,
        column_names_rot = 45,
        show_parent_dend_line = FALSE,
        row_dend_width = unit(25, "mm"))
```

```
## `use_raster` is automatically set to TRUE for a matrix with more than
## 2000 rows. You can control `use_raster` argument by explicitly setting
## TRUE/FALSE to it.
## 
## Set `ht_opt$message = FALSE` to turn off this message.
```

```
# plot average of each time point
avg <- sapply(seq(2, ncol(dds_vst.mat.var), 3), function(j) rowMeans(dds_vst.mat.var[, j+(-1:1)]))
colnames(avg) <- c("Cnt_1", "Cnt_2", "Cnt_3", "Cnt_10", "NaBut_1", "NaBut_2", "NaBut_3", "NaBut_10")
avg <- as.data.frame(avg)
avg$cluster <- unname(clust$cluster)
avg <- melt(avg)
```

```
## Warning in melt(avg): The melt generic in data.table has been passed a
## data.frame and will attempt to redirect to the relevant reshape2 method;
## please note that reshape2 is deprecated, and this redirection is now
## deprecated as well. To continue using melt methods from reshape2 while both
## libraries are attached, e.g. melt.list, you can prepend the namespace like
## reshape2::melt(avg). In the next version, this warning will become an error.
```

```
## Using cluster as id variables
```

```
avg$condition <- c(rep("Nt", nrow(avg)/2), rep("NaBut", nrow(avg)/2))
avg$time <- rep(c(1,2,3,10,1,2,3,10), each = nrow(avg)/8)
avg$gene <- rep(rownames(dds_vst.mat.var), times = 8)
avg <- unite(avg, unique_id, c(condition, gene), sep="_", remove = FALSE)

library(ggplot2)
library(ggpubr)

ggplot(avg, aes(x=time, y=value)) + 
  scale_color_manual(values = c("#006837", "#7f7f7f")) +
  scale_fill_manual(values = c("#006837", "#7f7f7f")) +
  theme_classic() +
  #geom_line(aes(group = unique_id, colour = condition), alpha = 0.05) +
  stat_summary(aes(y = value, colour = condition), fun = mean, geom = "line") +
  stat_summary(fun.data = mean_cl_boot,geom = "ribbon",size = 1, aes(fill = condition),alpha = 0.4) +
  facet_wrap(~ cluster, ncol=3, , scales="free") +
  scale_x_continuous(breaks=c(1,2,3,10)) +
  ylab("Normalized gene expression")
```

Gene ontology enrichments

```
library(clusterProfiler)
```

```
##
```

```
## Registered S3 method overwritten by 'ggtree':
##   method      from 
##   identify.gg ggfun
```

```
## clusterProfiler v4.2.2  For help: https://yulab-smu.top/biomedical-knowledge-mining-book/
## 
## If you use clusterProfiler in published research, please cite:
## T Wu, E Hu, S Xu, M Chen, P Guo, Z Dai, T Feng, L Zhou, W Tang, L Zhan, X Fu, S Liu, X Bo, and G Yu. clusterProfiler 4.0: A universal enrichment tool for interpreting omics data. The Innovation. 2021, 2(3):100141
```

```
## 
## Attaching package: 'clusterProfiler'
```

```
## The following object is masked from 'package:purrr':
## 
##     simplify
```

```
## The following object is masked from 'package:biomaRt':
## 
##     select
```

```
## The following object is masked from 'package:IRanges':
## 
##     slice
```

```
## The following object is masked from 'package:S4Vectors':
## 
##     rename
```

```
## The following object is masked from 'package:stats':
## 
##     filter
```

```
library(org.Hs.eg.db)
```

```
## Loading required package: AnnotationDbi
```

```
## 
## Attaching package: 'AnnotationDbi'
```

```
## The following object is masked from 'package:clusterProfiler':
## 
##     select
```

```
## The following object is masked from 'package:dplyr':
## 
##     select
```

```
##
```

```
df <- data.frame(cluster = clust$cluster, row.names = names(clust$cluster))
df <- split(df, f = df$cluster)
df <- lapply(df, function(x) row.names(x))

ck.bp <- compareCluster(geneCluster = df, 
                        fun = enrichGO,
                        keyType = "ENSEMBL",
                        OrgDb = org.Hs.eg.db,
                        ont = "BP",
                        readable = TRUE,
                        universe = rownames(dds_vst.mat),
                        qvalueCutoff = 0.05)
ck.bp <- simplify(ck.bp)
dotplot(ck.bp, showCategory=5, label_format = 100,) + ggtitle("Biological Process")
```

```
write.csv(ck.bp@compareClusterResult, "/Users/ronaldcutler/Library/CloudStorage//My\ Drive/Sidoli_Lab/Projects/Chromatin\ Decondensation/Analysis/Gencode/time-series_NaBut_vs_control_BP-enrichment.csv")

ck.mf <- compareCluster(geneCluster = df, 
                        fun = enrichGO,
                        keyType = "ENSEMBL",
                        OrgDb = org.Hs.eg.db,
                        ont = "MF",
                        readable = TRUE,
                        universe = rownames(dds_vst.mat),
                        qvalueCutoff = 0.05)
ck.mf <- simplify(ck.mf)
dotplot(ck.mf, showCategory=5, label_format = 50,) + ggtitle("Molecular Function")
```

```
write.csv(ck.mf@compareClusterResult, "/Users/ronaldcutler/Library/CloudStorage//My\ Drive/Sidoli_Lab/Projects/Chromatin\ Decondensation/Analysis/Gencode/time-series_NaBut_vs_control_MF-enrichment.csv")

ck.cc <- compareCluster(geneCluster = df, 
                        fun = enrichGO,
                        keyType = "ENSEMBL",
                        OrgDb = org.Hs.eg.db,
                        ont = "CC",
                        readable = TRUE,
                        universe = rownames(dds_vst.mat),
                        qvalueCutoff = 0.05)
ck.cc <- simplify(ck.cc)
dotplot(ck.cc, showCategory=5, label_format = 50,) + ggtitle("Cellular Component")
```

```
write.csv(ck.cc@compareClusterResult, "/Users/ronaldcutler/Library/CloudStorage//My\ Drive/Sidoli_Lab/Projects/Chromatin\ Decondensation/Analysis/Gencode/time-series_NaBut_vs_control_CC-enrichment.csv")
```

Cell type signature enrichment

```
library(msigdbr)
# get cell type signature gene set
C8_t2g <- msigdbr(species = "Homo sapiens", category = "C8") %>% 
  dplyr::select(gs_name, ensembl_gene)

ck.c8 <- compareCluster(geneCluster = df, 
                        fun = enricher,
                        TERM2GENE = C8_t2g,
                        universe = rownames(dds_vst.mat),
                        qvalueCutoff = 0.05)

dotplot(ck.c8, showCategory=5, label_format = function(x) stringr::str_wrap(x, width=40)) + ggtitle("MSigDB C8 Cell Type Signatures")
```

```
# unable to just subset liver only 
#ck.c8.liver <- ck.c8@compareClusterResult$ID[grep("LIVER", ck.c8@compareClusterResult$ID)
#dotplot(ck.c8, showCategory=ck.c8.liver, label_format = function(x) stringr::str_wrap(x, width=40)) + ggtitle("Cell Type Signatures")
```

Liver curated gene sets enrichment

```
# get cell type signature gene set
C2_t2g <- msigdbr(species = "Homo sapiens", category = "C2") %>% 
  dplyr::select(gs_name, ensembl_gene)
C2_t2g <- C2_t2g[grep("LIVER", C2_t2g$gs_name),]
  
ck.c2 <- compareCluster(geneCluster = df, 
                        fun = enricher,
                        TERM2GENE = C2_t2g,
                        universe = rownames(dds_vst.mat),
                        qvalueCutoff = 0.05)

dotplot(ck.c2, showCategory=5, label_format = function(x) stringr::str_wrap(x, width=40)) + ggtitle("MSigDB C2 Curate Signature for Liver")
```
